# SLC44A1 deficiency impedes myelin development in the central nervous system

**DOI:** 10.1101/2025.04.08.647819

**Authors:** Qiang Chen, Xiang Chen, Zhenghao Li, Qi Shao, Hao Huang, Wenhao Zhou, Mengsheng Qiu, Zhida Su, Peng Liu, Cheng He

**Author notes:** These authors contributed equally (Qiang Chen, Xiang Chen, Zhenghao Li). Correspondence (P.L.), (C.H.).

## Abstract

Axon-wrapping myelin sheath is essential for the proper functioning of the central nervous system (CNS). Defects in myelination occur in various neurodevelopmental disorders. Human deficiency of the solute carrier 44A1 (SLC44A1) causes a new type of childhood-onset neurodegeneration with leukoencephalopathy. Currently, there is no effective treatment for this devastating disease, and little is known about its pathological mechanisms. In this study we identified that SLC44A1 is enriched in oligodendrocytes and required for myelin development in the CNS of zebrafish and rodents, which is crucial for normal motor coordination and cognitive function. Defective oligodendroglial maturation and myelinogenesis can be visualized by *in vivo* time-lapse imaging in the spinal cord of zebrafish with *Slc44a1b* deficiency. Mechanistically, SLC44A1 deficiency disrupts the expression of genes involved in phosphatidylcholine production pathway, and subsequently inhibiting phospholipid biosynthesis and disturbing the lipid composition of myelin sheaths. More importantly, supplementation with citicoline, a natural choline metabolite, restores developmental myelination in SLC44A1-deficient animals. These findings demonstrate that SLC44A1 is essential for CNS myelination and citicoline supplementation represents a potential therapeutic strategy to address the developmental hypomyelination.

## 1. Introduction

Myelination by oligodendrocytes (OLs) enables saltatory propagation of action potentials and provides long-term trophic support for neuronal axons, thereby maintaining neural integrity in both white and grey matter of the central nervous system (CNS).^1,2^ Impairments in myelination disrupt the transmission of nerve impulses and the integrity of functional connectivity, resulting in cognitive and motor deficits associated with a range of neurodevelopmental disorders.^3–6^ However, the molecular barriers that impede myelin development remain poorly understood.

Choline is a quaternary ammonium cation and water-soluble compound essential for cellular functions.^7–9^ It serves as a precursor for the neurotransmitter acetylcholine,^10–13^ essential for learning and memory, and contributes to phospholipid synthesis,^14–17^ which is necessary for cellular membrane structure and function. Choline also provides one-carbon units for numerous biochemical reactions.^18–20^ Due to the fact that choline is a hydrophilic cationic molecule that does not freely permeate cell membranes, its uptake depends on carrier-mediated transport.^21^ Solute carrier 44A1 (SLC44A1), also known as choline transporter-like protein 1, is considered as a universal choline transporter across both the plasma and mitochondrial membrane.^22,23^ SLC44A1 has been implicated in various diseases, including motor disorder^24^ and tumor development^25^. Polymorphisms in SLC44A1 are identified to associate with cognitive decline in children^26,27^ Moreover, a genetical study reports that the homozygous frameshift mutations in SLC44A1 cause a new type of childhood-onset neurodegeneration with clinical features including progressive ataxia, tremor, cognitive decline, optic atrophy, dysphagia, and dysarthria, and brain MRI reveals cerebellar atrophy and leukoencephalopathy.^28^ Currently, there is no effective treatment for this devastating neurodevelopmental disorder, and the underlying molecular mechanism has not been investigated.

In the present study, we demonstrated that SLC44A1 is enriched in OLs and required for the developmental myelination in zebrafish and rodents. SLC44A1 deficiency impairs the biosynthesis of phosphatidylcholine (PC) and disrupts the phospholipid composition of myelin membrane. Importantly, citicoline, a choline metabolic product, alleviates the defects of myelin formation in SLC44A1 mutant animals, providing new potential strategies for clinical therapies.

## 2. Results

### 2.1. SLC44A1 is enriched in OLs and correlates with myelination

To elucidate the expression pattern of SLC44A1 in the CNS of mice, by using publicly available datasets (GSE176063 and GSE198832),^29,30^ we first performed clustering analysis on the corresponding single-cell RNA sequencing data from murine CNS cells. We identified nine distinct cell clusters with specific gene expression profiles (**Figure 1A, B**). Interestingly, we found that *Slc44a1* was highly expressed in *Mbp* and *Mag*-enriched mature OL clusters (Figure 1C; Figure S1A). Gene violin plots also revealed that *Slc44a1* was highly enriched in mature OLs, with minimal expression in other cells in the CNS, such as oligodendrocyte precursor cells (OPCs), neurons, microglia and astrocytes (Figure 1D).

**Figure 1.**
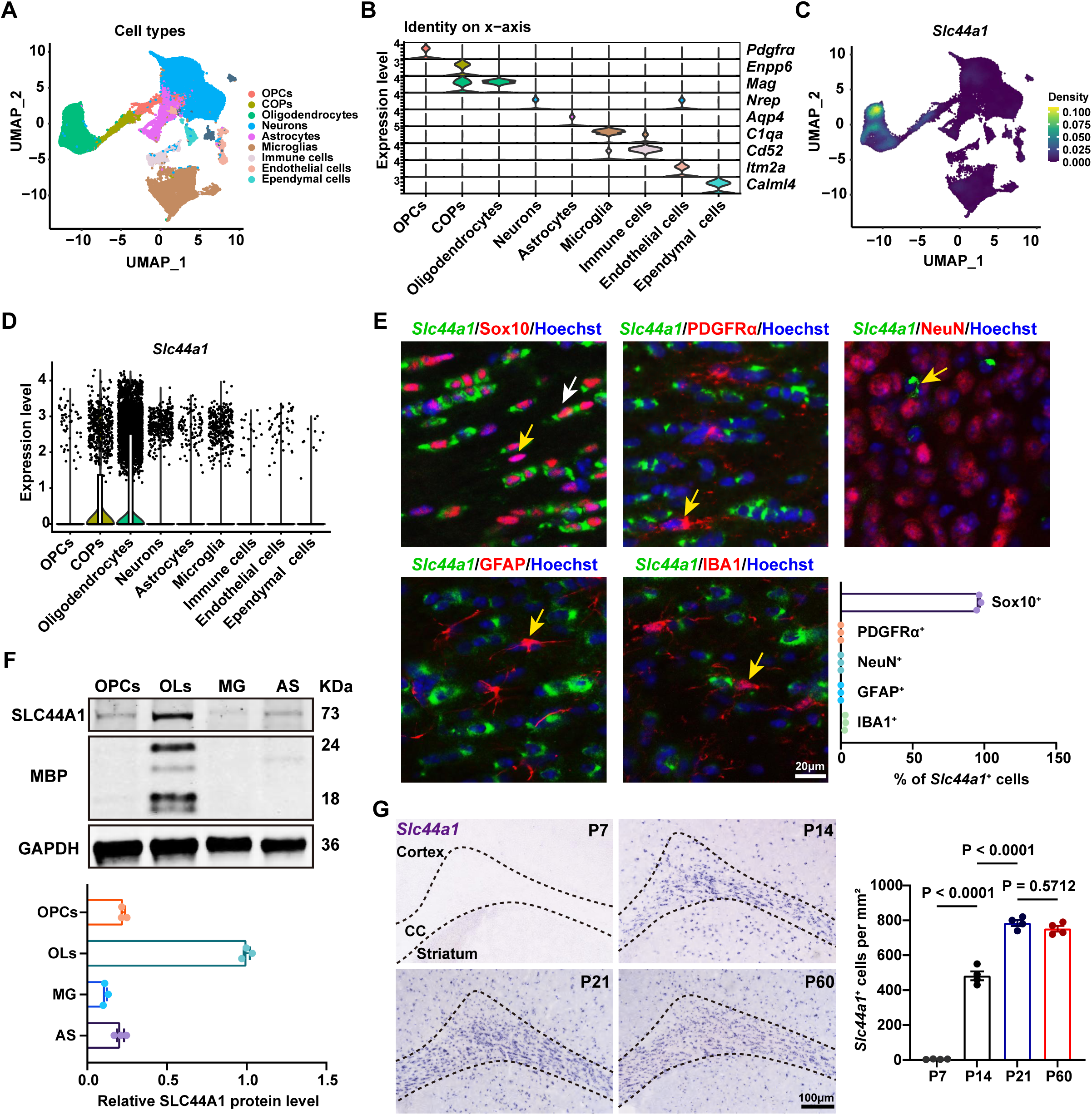
SLC44A1 is enriched in OLs. (**A**) Uniform manifold approximation and projection (UMAP) plots identifying OPCs, COPs (differentiation-directed OPCs), OLs, neurons, astrocytes, microglia, immune cells, endothelial cells, and ependymal cells in mouse brain. (**B**) Violin plot showing the distribution of expression levels of well-known representative cell-type-enriched marker genes across all 9 cell types. (**C**) UMAP plots showing the expression of *Slc44a1* mRNA. (**D**) Violin plots showing the expression of *Slc44a1* mRNA in each cell type. (**E**) *In situ* hybridization for *Slc44a1* (green) and immunostaining for Sox10, PDGFRα, GFAP, or IBA1 (red) in the CC of wildtype mice at P21, and immunostaining for NeuN (red) in the cortex of wildtype mice at P21. White arrow indicates *Slc44a1*^+^ Marker^+^ cells. Yellow arrows indicate Marker^+^ cells without *Slc44a1* expression. Right corner, percentage of *Slc44a1*^+^ and Marker^+^ cells among *Slc44a1*^+^ cells (n = 3 mice). (**F**) Western blots and quantification of SLC44A1 protein level in primary rat glia cells (n = 3 independent experiments). (**G**) *In situ* hybridization showing expression of *Slc44a1* (purple) at the indicated time points and quantification of *Slc44a1*^+^ cell numbers in the CC of wildtype mice (n = 3 mice).

To further determine the cellular localization of SLC44A1 in the brain, we generated an *in situ* RNA probe specific for *Slc44a1* and performed co-localization staining with several neural cell markers. Consistent with the single-cell RNA-seq data, the results showed that *Slc44a1* was abundantly expressed in Sox10^+^ OLs, but nearly undetectable in PDGFRα^+^ OPCs in the corpus callosum (CC) at postnatal day 21 (P21) (Figure 1E). Furthermore, *Slc44a1* was minimally expressed in NeuN^+^ neurons, GFAP^+^ astrocytes, and IBA1^+^ microglia (Figure 1E). In culture, SLC44A1 expression was predominantly detected in mature OLs via Western blotting (Figure 1F). Consistently, MBP^+^ mature OLs were strongly immunolabelled for SLC44A1 (Figure S1B).

*Slc44a1* expression in the white matter also exhibited dynamic change. *Slc44a1*^+^ cells were scarce around P7 and more detectable at P14, reached a peak at P21, and persisted into adulthood (Figure 1G), which correlates with the timeline of brain myelination.^31^ Furthermore, the majority of *Slc44a1*^+^ cells were Sox10^+^ OLs throughout development (Figure S1C, D). The dynamic expression pattern of *Slc44a1* in the brain was highly consistent with the results from the single-cell data (Figure S1E). Together, the temporal and spatial expression characteristics suggest that SLC44A1 may be associated with OL maturation and myelination.

### 2.2. SLC44A1 is critical for developmental myelination

To investigate the function of SLC44A1 in the CNS myelination, we initially employed the CRISPR-Cas9 technology to generate conditional *Slc44a1* knockout mice with a deletion of the eighth exon. By crossing mice carrying floxed *Slc44a1* alleles with the *Olig1*-Cre line,^32^ we successfully obtained the conditional mutant mice in which *Slc44a1* is selectively ablated in OL lineage (*Slc44a1*^flox/flox^; *Olig1*-Cre^+/-^, hereafter referred to as *Slc44a1* cKO) (**Figure 2**A). Consistently, conditional mutation results in a significant reduction in the number of *Slc44a1*^+^ cells in the CC (Figure 2B).

**Figure 2.**
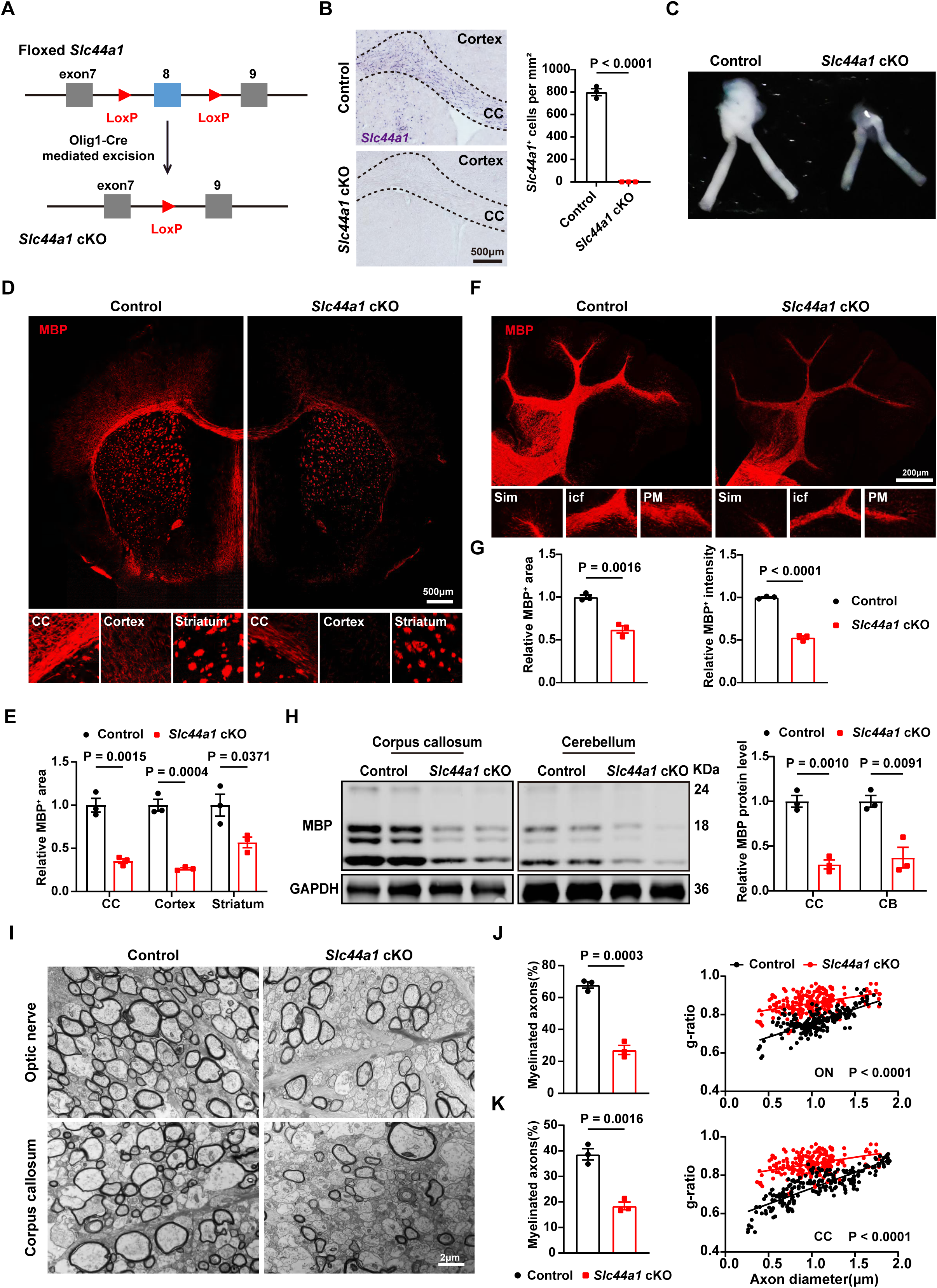
Deletion of SLC44A1 in the OL lineage exhibits hypomyelination during development. (**A**) Diagram depicting generation of *Slc44a1* cKO mice. (**B**) Left, *in situ* hybridization for *Slc44a1* in the cortex and CC of control (*Slc44a1^flox/flox^*) and *Slc44a1* cKO (*Slc44a1^flox/flox^; Olig1-Cre^+/-^*) mice at P21. Right, quantification of *Slc44a1*^+^ cell number in the CC (n = 3 mice). (**C**) Representative images of ON from control and *Slc44a1* cKO mice at P21. (**D**) MBP immunostaining in multiple brain regions of control and *Slc44a1* cKO mice at P21. (**E**) Quantification of relative MBP^+^area in the cortex, CC, and striatum of control and *Slc44a1* cKO at P21 (n = 3 mice). (**F**) MBP immunostaining in the cerebellum of control and *Slc44a1* cKO mice at P21. Sim, simple lobule; icf, intercrural fissur; PM, paramedian lobule. (**G**) Quantification of relative MBP^+^ area and intensity in the cerebellum of control and *Slc44a1* cKO mice at P21 (n = 3 mice). (**H**) Western blots and quantification of MBP protein level in CC and cerebellum of control and *Slc44a1* cKO mice at P21 (n = 3 mice). (**I**) Electron micrographs of ON and CC of control and *Slc44a1* cKO mice at P21. (**J**, **K**) Left, percentage of myelinated axons in the ON and CC of control and *Slc44a1* cKO mice at P21 (n = 3 mice). Right, scatter plot of g-ratio from the ON and CC of P21 control and *Slc44a1* cKO mice (>200 axons from each genotype; n = 3 mice, each point representing one axon).

The myelinated optic nerve (ON) of *Slc44a1* cKO mice appeared translucent at P21, indicating a myelin deficiency (Figure 2C). Immunostaining study confirmed that MBP expression in different CNS regions of P21 *Slc44a1* cKO mice, including the CC, cerebral cortex, striatum (Figure 2D, E), and cerebellum (Figure 2F, G), were significantly lower than that in controls. Similarly, mRNA *in situ* hybridization results showed that the expression levels of *Mbp* (Figure S2A, B) and *Plp1* (Figure S2C, D) were also markedly decreased in the CC, cerebral cortex, and striatum of *Slc44a1* cKO mice during development. In addition, western blotting revealed that the protein levels of MBP in the CC and cerebellum of *Slc44a1* cKO mice were substantially lower than that in control mice (Figure 2H).

Next, we employed transmission electron microscopy (TEM) to examine the ultrastructural changes of myelin in white matter tracts such as the ON and CC. The degree of axonal myelination in *Slc44a1* cKO mice was severely diminished, evidenced by a reduced percentage of myelinated axons and a decrease in myelin thickness within the ON and CC of P21 *Slc44a1* cKO mice compared to control mice (Figure 2I-K). Thus, conditional deletion of *Slc44a1* in OLs impedes developmental myelination in mice.

### 2.3. SLC44A1 deletion impairs OL maturation and myelinogenesis

To investigate whether the hypomyelination phenotype observed in *Slc44a1* cKO mice is due to defects in OL development, we initially assessed OPC numbers by *in situ* staining of *Pdgfra* at P7. The number of *Pdgfra*^+^ cells in *Slc44a1* cKO mice was comparable to that in controls (Figure S3A). Similar result was also achieved by PDGFRα immunostaining in the CC at P21 (**Figure 3**A). Furthermore, Ki67 and BrdU immunostaining revealed comparable OPC proliferative rates in *Slc44a1* cKO mice and in cultures with *Slc44a1* knock-down by lentiviral interference as compared to the controls (Figure S3B-D).

**Figure 3.**
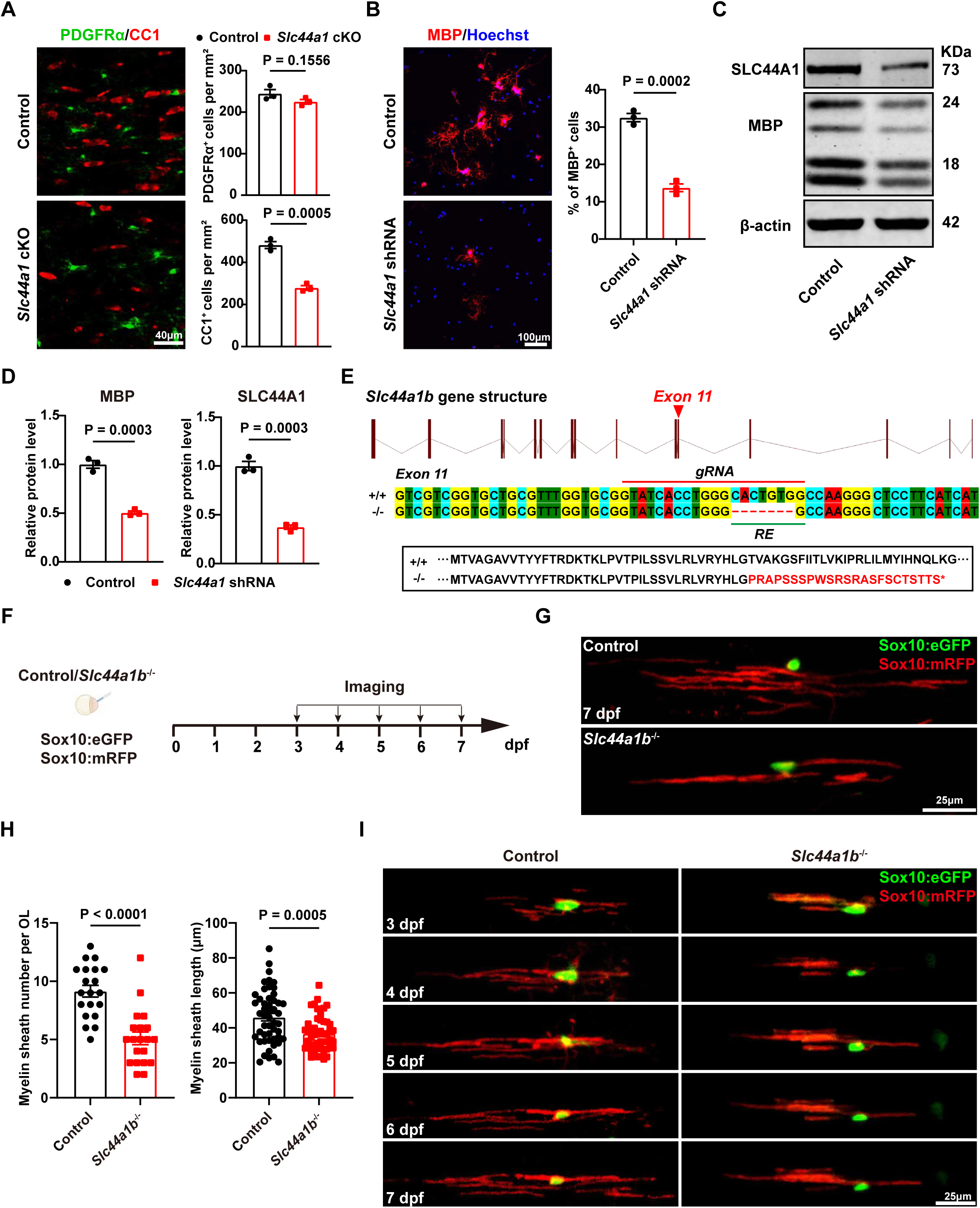
SLC44A1 loss-of-function impairs OL maturation and myelinogenesis. (**A**) Left, immunostaining for CC1 and PDGFRα in the CC of control and *Slc44a1* cKO mice at P21. Right, quantification of CC1^+^ and PDGFRα^+^ cells (n = 3 mice). (**B**) Left, immunostaining for MBP in control and *Slc44a1* shRNA-treated OLs. Right, quantification of MBP^+^ cells (n = 3 independent experiments). (**C**) Western blots of MBP and SLC44A1 in control and *Slc44a1* shRNA-treated OLs. (**D**) Quantification of MBP and SLC44A1 protein levels in control and *Slc44a1* shRNA-treated OLs (n = 3 independent experiments). (**E**) Top, *Slc44a1b* gene structure composed of 15 exons. Red arrowhead marks the location of the mutation in exon 11. Middle, Wild-type and mutant nucleotide sequences spanning the mutagenesis site. The *gRNA* target site (red line) is labeled. Bottom, Amino acid sequence indicating that the *Slc44a1b* mutation results in shift in the open reading frame leading to downstream coding for a premature stop codon (*). (**F**) Diagram of single OL mosaically labeled by Sox10:eGFP and Sox10:mRFP transient expression in controls and *Slc44a1b*^-/-^ mutants. Timeline indicating the imaging procedures. (**G**) Typical case of single OL and its myelin segments labelled from control and *Slc44a1b*^-/-^ larvae at 7 dpf. (**H**) Statistics of the number and length of myelin sheath segments from an individual OLs from control and *Slc44a1b*^-/-^ larvae at 7 dpf. (**I**) Representative time-lapse images of an individual OLs from control and *Slc44a1b*^-/-^ larvae, across various time points (3 - 7dpf).

Given the abundant expression in mature OLs, we postulated that SLC44A1 is required for OL maturation. As hypothesized, there was a significant reduction in the number of CC1^+^ mature OLs in the CC of *Slc44a1* cKO mice when compared to control mice (Figure 3A). The number of MBP^+^ cells and MBP protein expressions were markedly decreased following *Slc44a1* knockdown in cultured OLs (Figure 3B-D). OL survival appeared to be unaffected by the *Slc44a1* conditional knockout, as confirmed by cleaved caspase-3 immunostaining in the CC (Figure S3E). These findings suggest that SLC44A1 is required for OL maturation.

To further examine the myelin-producing capacity of OLs *in vivo*, we generated frame-shift mutant zebrafish by CRISPR-Cas9 technology with 7 bases ablation in exon 11 of *Slc44a1b* gene (*Slc44a1b*^-/-^) (Figure 3E). Gross morphologies e.g. the body length and eye area of embryos were comparable between *Slc44a1b* mutants and controls (Figure S4A-D). We then preformed long-term imaging of OLs in the spinal cord (SC) of Tg(Olig2:eGFP) and Tg(Sox10:mRFP) lines, which labeled the soma of OL lineage cells and myelin sheaths, respectively, from 3 - 5 days post-fertilization (dpf) with 24 h intervals (Figure S4A, E). No significant difference in the number of OL linage cells was found in the dorsal SC (dSC), which originated from the ventral SC, at 3 dpf (the early stage of oligodendrogenesis in the SC, with most of the GFP positive cells being OPCs). However, a markable decline at 4 - 5dpf (the stage of myelination) was observed in *Slc44a1b* mutants compared with controls (Figure S4E, F). Moreover, the area and intensity of mRFP, which labeled the myelin sheath segments, were reduced dramatically in the dSC from 3 to 5 dpf in *Slc44a1b* mutants (Figure S4E, G, H). These results indicate that SLC44A1 was required for OL maturation and myelination, in keeping with our findings in rodents.

To visualize the myelinating procession of single OL, we further conducted time-lapse observation of an individual OL mosaically labeled by Sox10:eGFP and Sox10:mRFP transient expression, which marked the soma and myelin sheaths of an OL respectively, in zebrafish SC from 3 to 7 dpf with 24 h intervals (Figure 3F). It was found that both the length and number of myelin sheath segments formed by individual OL decreased significantly in *Slc44a1b* mutants at 7 dpf (Figure 3G, 3H), and the dynamic myelinating process of OLs was also impaired by *Slc44a1b* deficiency from 3 to 7 dpf (Figure 3I). Thus, these results indicate that SLC44A1 is essential for the myelin production of OLs.

### 2.4. Oligodendroglial SLC44A1 deficiency impairs motor coordination and cognitive function

To determine whether the deficits in developmental myelination persists into adulthood in *Slc44a1* cKO mice, we evaluated the extent of myelination in the adult brains. Black gold staining revealed a substantial reduction in myelin density throughout the brains of adult *Slc44a1* cKO mice, particularly in the cortex, CC and striatum (**Figure 4**A, B). Immunostaining for myelin proteins, including MBP and PLP, revealed a marked decrease in the area and intensity of MBP^+^ and PLP^+^ fluorescence in the whole brain of adult *Slc44a1* cKO mice (Figure 4C, D; Figure S5A-D). Furthermore, we also observed hypomyelination in the cerebellum of adult *Slc44a1* cKO mice through MBP immunostaining (Figure 4E, F). Collectively, these findings suggest that myelination deficits are sustained in the brains of adult *Slc44a1* cKO mice.

**Figure 4.**
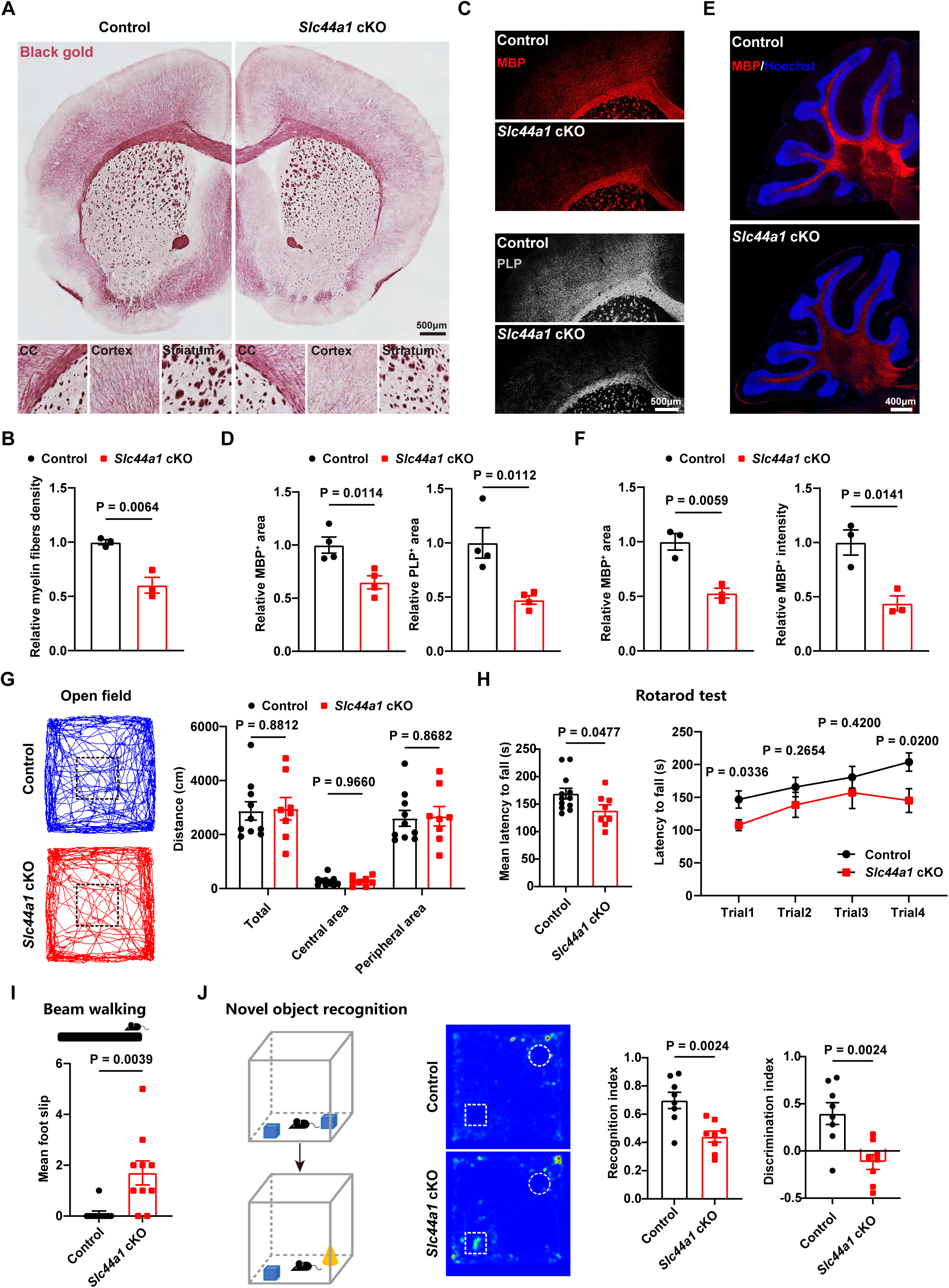
The deficiency of SLC44A1 in OLs causes impairment of motor coordination and cognitive function. (**A**) Black gold myelin staining in multiple brain regions of control and *Slc44a1* cKO mice at P60. (**B**) Quantification of relative myelin fibers density in the whole brain of control and *Slc44a1* cKO at P60 (n = 3 mice). (**C**) Immunostaining for MBP and PLP in cortex and CC of control and *Slc44a1* cKO mice at P60. (**D**) Quantification of relative MBP^+^ and PLP^+^ area in cortex and CC of control and *Slc44a1* cKO mice at P60 (n = 4 mice). (**E**) Immunostaining for MBP in the cerebellum of control and *Slc44a1* cKO mice at P60. (**F**) Quantification of relative MBP^+^ area and intensity in the cerebellum of control and *Slc44a1* cKO mice at P60 (n = 3 mice). (**G**) Left, open field trajectory maps of control and *Slc44a1* cKO mice at P60. Right, quantification of the distance of control and *Slc44a1* cKO mice in the entire area, center area, and peripheral area of the open field (control: n= 10 mice; *Slc44a1* cKO: n= 10 mice). (**H**) Left, quantification of the mean latency to fall from the spinning drum in four independent rotarod tests. Right, quantification of the latency to fall from the spinning drum in each rotarod test (control: n= 12 mice; *Slc44a1* cKO: n = 8 mice). (**I**) Top, diagram of beam walking test. Bottom, quantification of mean foot slips of control and *Slc44a1* cKO mice (control: n=10 mice; *Slc44a1* cKO: n= 10 mice). (**J**) Left, diagram of novel object recognition test. Center, trajectory heatmaps of control and *Slc44a1* cKO mice at P60. Right, quantification of the recognition index and discrimination index of control and *Slc44a1* cKO mice (control: n = 10 mice; *Slc44a1* cKO: n = 10 mice).

It is widely recognized that hypomyelination significantly disrupts the proper functions of CNS, especially for the modulation of fine motor skills and cognitive processes.^33,34^ Accordingly, we examined whether myelination deficits in adult *Slc44a1* cKO mice lead to behavioral abnormalities. In the open field test, there were no significant differences in the total travel distance between *Slc44a1* cKO mice and control mice, suggesting unaffected spontaneous motor function of *Slc44a1* cKO mice (Figure 4G). Furthermore, the distance traveled in both the center and periphery of the open field remained unchanged, indicating that *Slc44a1* cKO mice do not exhibit an anxiety-like phenotype (Figure 4G). The rotarod test and beam walking test were used to assess motor coordination and balance. In four consecutive independent rotarod tests, the average latency to fall off the rod was significantly reduced in *Slc44a1* cKO mice, especially in the first and fourth trials (Figure 4H). The beam walking test revealed a significant increase in the number of times that *Slc44a1* cKO mice slipped with their hind limbs during the fixed-length walk on the beam (Figure 4I). In addition, in the novel object recognition test, a commonly employed method for evaluating recognition memory in mice, showed that *Slc44a1* mutants displayed a lack of preference for exploring the novel object, accompanied by a notable decline in both recognition and discrimination indices (Figure 4J). Taken together, these results suggest that adult *Slc44a1* cKO mice exhibit impaired motor coordination, balance, and cognitive function.

### 2.5. SLC44A1deficiency disrupts gene expression involved in myelin formation and choline metabolism

To elucidate the mechanisms underlying the impaired myelin formation following *Slc44a1* deficiency, we carried out RNA sequencing (RNA-seq) analysis on primary cultured OLs treated with *Slc44a1* interfering lentivirus. In comparison to the control group, the expression of 1,430 genes was significantly altered by *Slc44a1* knockdown (**Figure 5**A). Notably, genes involved in the regulation of myelin formation, including *Mbp*, *Mag*, *Mog*, *Cnp*, *Aspa*, and *Enpp6*, exhibited significant downregulation in the *Slc44a1* shRNA-treated group (log_2_(fold-change) ≤ -0.5, P ≤ 0.01). Gene Set Enrichment Analysis (GSEA) revealed that the downregulated genes in *Slc44a1* knockdown OLs were predominantly enriched in gene sets related to OL maturation and myelination (Figure 5B). Similarly, Gene Ontology (GO) enrichment analysis also showed that significantly downregulated genes were mainly enriched in biological processes associated with myelination and myelin sheath (Figure 5C), which was consistent with the myelin developmental phenotype observed in *Slc44a1* cKO mice. Quantitative PCR (qPCR) analysis further confirmed the reduction of myelination-related genes expression (Figure 5D).

**Figure 5.**
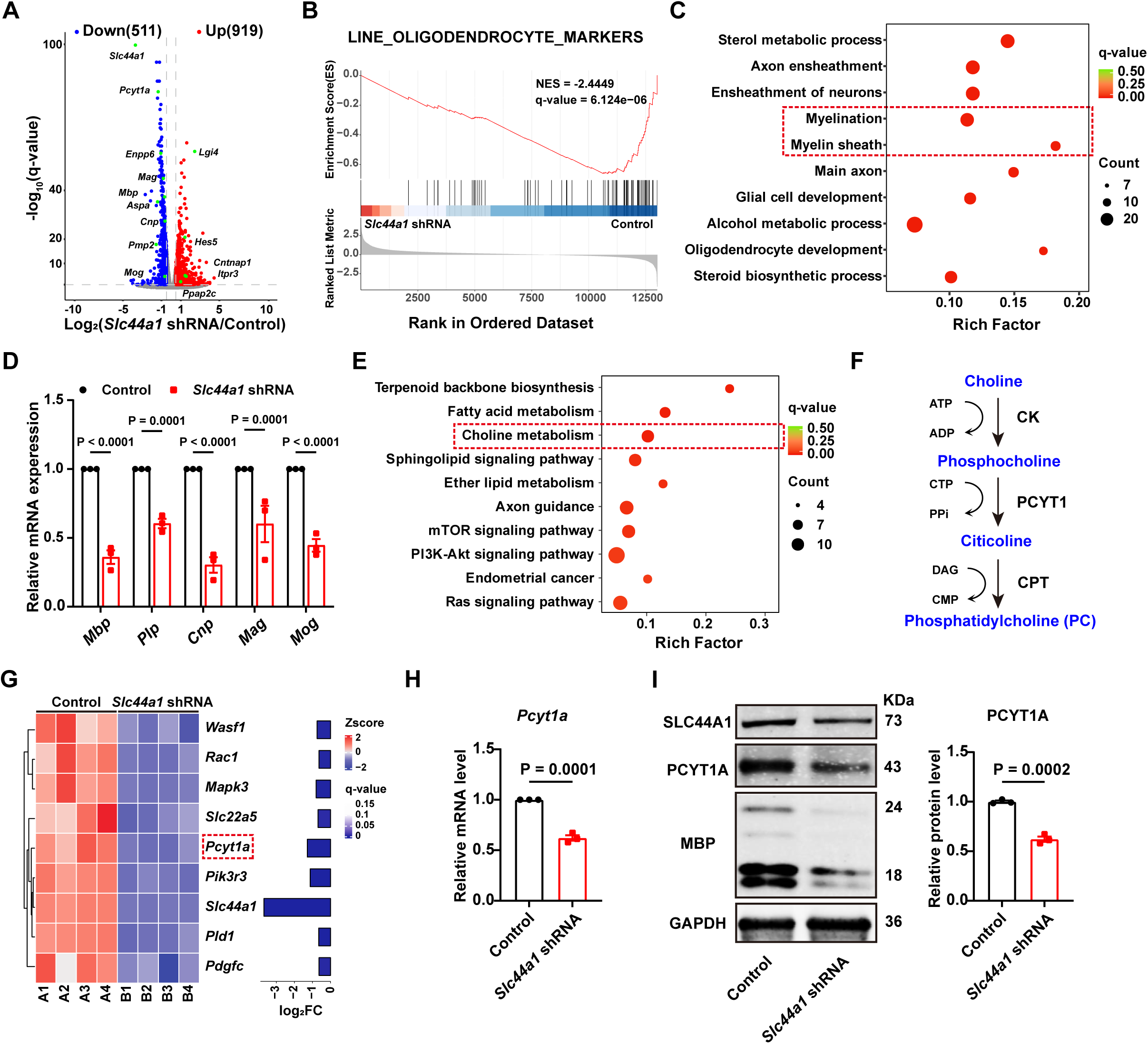
The absence of SLC44A1 disrupts the transcriptome associated with myelinogenesis and choline metabolism. (**A**) Volcano plots of gene expression differences between control and *Slc44a1* shRNA-treated OLs (n = 4 experiments analyzed by DESeq2; |log_2_(fold-change)| ≥ 0.5, P ≤ 0.01). (**B**) GSEA plot showing OL signature genes downregulated in the *Slc44a1* shRNA-treated OLs. NES, net enrichment score. (**C**) GO enrichment analysis of downregulated genes in *Slc44a1* shRNA-treated OLs. (**D**) qPCR analysis of myelination-associated genes in control and *Slc44a1* shRNA-treated OLs (n = 3 experiments). (**E**) KEGG pathway enrichment analysis of downregulated genes in *Slc44a1* shRNA-treated OLs. (**F**) Schematics of choline metabolism pathway. ATP, adenosine triphosphate; ADP, adenosine diphosphate; CTP, cytidine triphosphate; PPi, pyrophosphate; DAG, diacylglycerol; CMP, cytidine monophosphate; CK, choline kinase; PCYT1A, phosphate cytidylyltransferase 1A; CPT, choline phosphate transferase. (**G**) Left, heatmap of downregulated differential genes in choline metabolism in *Slc44a1* shRNA-treated OLs (n = 4 experiments). Right, fold change of differential genes. (**H**) qPCR analysis of *Pcyt1a* mRNA level in control and *Slc44a1* shRNA-treated OLs (n = 3 experiments). (**I**) Western blots and quantification of PCYT1A protein level in control and *Slc44a1* shRNA-treated OLs (n = 3 experiments).

Additionally, Kyoto Encyclopedia of Genes and Genomes (KEGG) pathway enrichment analysis showed that the majority of downregulated genes were significantly enriched in metabolism-related pathways, particularly the choline metabolic pathway (Figure 5E). Choline metabolism plays a crucial role in the synthesis of PC, which is essential for cell membrane growth and maintenance^35^ (Figure 5F). The heatmap analysis revealed that, in addition to *Slc44a1*, *phosphatidylcholine cysteine lyase type 1a* (*Pcyt1a*) was the most significantly downregulated choline metabolic gene following the knockdown of *Slc44a1* (Figure 5G). The PCYT1A protein, encoded by the *Pcyt1a* gene, acts as the primary rate-limiting enzyme in the choline metabolism, especially in PC synthesis.^36^ Both qPCR and western blotting confirmed a notable reduction in the expression of *Pcyt1a* mRNA and PCYT1A protein in OLs following SLC44A1 knockdown (Figure 5H, I). These data suggest that SLC44A1 may facilitate myelination through the modulation of the choline metabolic pathway.

### 2.6. SLC44A1 is required for choline metabolism in OLs and PC synthesis of myelin

To gain further insights into the role of SLC44A1 in choline metabolism of OLs, we performed the metabolomics analysis (**Figure 6**A). Volcano plot analysis showed that 86 metabolites exhibited significantly altered levels following the knockdown of SLC44A1, with 72 of these being downregulated (Figure 6B). The KEGG pathway analysis of the downregulated metabolites revealed an enrichment for the choline metabolic pathway (Figure 6C). Notably, among the metabolites downregulated in the choline metabolic pathway, the most significant changes were observed in citicoline which is the direct upstream metabolite for PC synthesis and catalytically formed by PCYT1A from phosphocholine (Figure 5F; Figure 6D).

**Figure 6.**
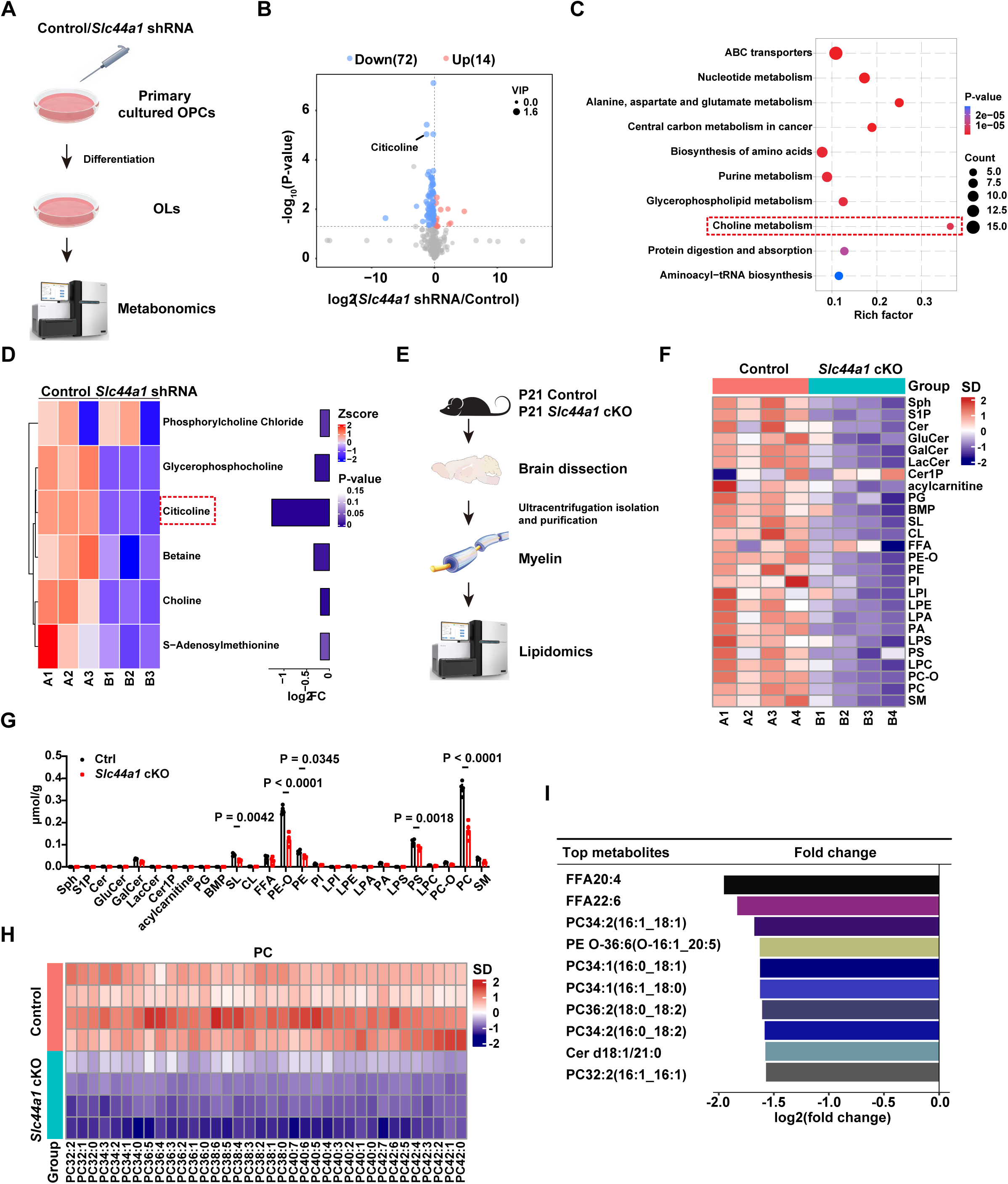
SLC44A1 loss-of-function impairs choline metabolism in OLs and phospholipid biosynthesis of myelin. (**A**) Schematic diagram of the Metabolomics workflow for control and Slc44a1 shRNA-treated OLs. (**B**) Volcano plots of the metabolite differences between control and *Slc44a1* shRNA-treated OLs (n = 3 experiments; VIP > 1, P < 0.05). VIP, variable importance in the projection. (**C**) KEGG pathway enrichment analysis of downregulated metabolites from control and *Slc44a1* shRNA-treated OLs. (**D**) Left, heatmap of downregulated metabolites in choline metabolism from control and *Slc44a1* shRNA-treated OLs (n = 3 experiments). Right, fold change of differential metabolites. (**E**) Schematic diagram of the lipidomics workflow for myelin components. (**F**) Heatmap of lipid components in myelin from control and *Slc44a1* cKO mice (n = 3 mice). (**G**) Quantification of lipid components in myelin from control and *Slc44a1* cKO mice (n = 3 mice). (**H**) Heatmap plotting the compositions of PC in myelin from control and *Slc44a1* cKO mice (n = 3 mice). (**I**) Top 10 of differential lipid components in myelin of *Slc44a1* cKO mice relative to controls (n = 3 mice).

To further understand the impact of SLC44A1 deficiency on PC synthesis of myelin, we extracted and purified myelin sheaths from the whole brains of control and *Slc44a1* cKO mice at P21 and performed lipidomics analysis (Figure 6E). Heatmap analysis of the lipid constituents in myelin sheaths showed a significant reduction in the majority of these lipid components in *Slc44a1* cKO mice when compared to control mice (Figure 6F). Significant differences were observed for sulfatides, alkyl phosphatidylethanolamines, phosphatidylethanolamines (PE), phosphatidylserines, and PC with the most pronounced decrease (Figure 6G). The PC composition analysis revealed that PC with varying chain lengths and degrees of unsaturation were reduced in the myelin sheaths of *Slc44a1* cKO mice (Figure 6H). Indeed, the top 10 lipids with the most significant declines exhibited the highest proportion of PCs in the myelin sheaths (Figure 6I). Collectively, these observations suggest that SLC44A1 is required for the synthesis of PC, a major lipid component of myelin sheaths, which is largely mediated by the regulation of citicoline.

### 2.7. Citicoline supplementation improves the myelination deficits caused by SLC44A1 deficiency

Next, we asked whether supplementation of citicoline could rescue developmental hypomyelination in *Slc44a1* mutant animals. First, citicoline was administrated to Tg(MBPa:eGFP-CAAX) transgenic zebrafish, which labeled myelin sheaths by GFP-CAAX, from 1 to 7 dpf and the extent of myelination was assessed by confocal imaging at 5 dpf and TEM at 7 dpf (**Figure 7**A). Supplementation of citicoline did not affect the overall morphology of control and mutant zebrafish larvae, as indicated by the body length and eye area (Figure 7B, C). Confocal imaging showed that citicoline did not influence the myelination patterns in controls, whereas it markedly ameliorated the myelination defects in mutants, as measured by MBP^+^ area and intensity in the dorsal and ventral SC (Figure 7D, E). Consistent with these findings, TEM analysis showed that citicoline significantly increased the density of myelinated axons in the dorsal and ventral SC in *Slc44a1* mutants (Figure 7F-H).

**Figure 7.**
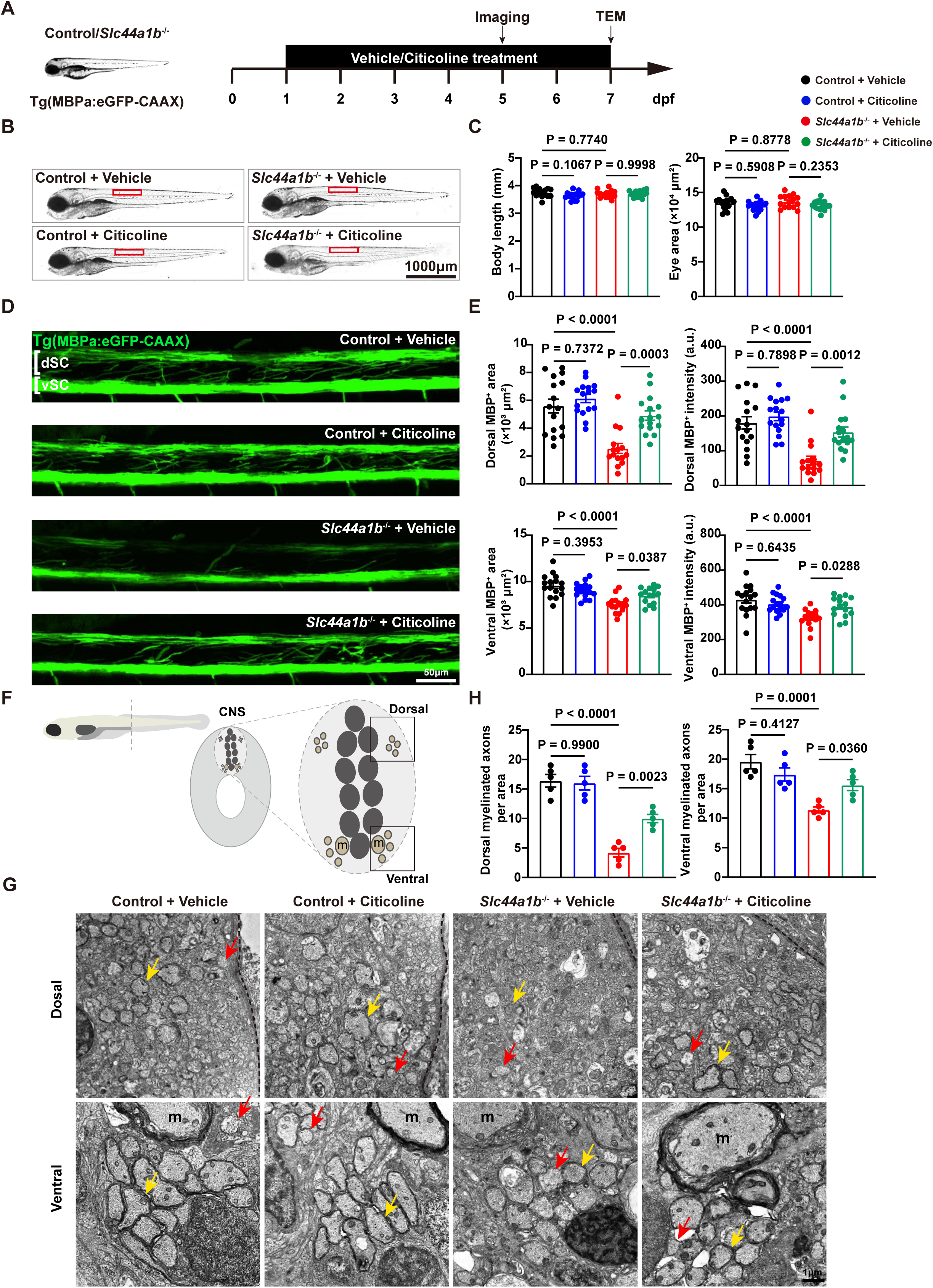
Citicoline supplementation rescues the developmental myelination deficits in *Slc44a1b* mutant fish. (**A**) Schematic diagram displaying the time course for citicoline treatment, imaging, and TEM analysis. TEM, transmission electron microscope. (**B**) Bright field imaging shows the normal morphology and the area (7 - 12 somites) being analyzed in Figure 7D. (**C**) Measurements of larval length and eye area (n ≥ 14 fish). (**D**) The fluorescent images of Tg(MBPa:eGFP-CAAX) transgenic line by control with vehicle, control with citicoline, *Slc44a1b*^-/-^ with vehicle, and *Slc44a1b*^-/-^ with citicoline. (**E**) Quantification of MBP^+^ area and intensity in the dorsal and ventral SC (n ≥ 14 fish). (**F**) Schematic of the transverse section of a 7 dpf larval zebrafish at the level of the urogenital opening. Inset, transverse section of the SC at the same level. Myelinated (yellow) axons are located in the dorsal and ventral spinal tracts of the SC (CNS). m, Mauthner axons. (**G**) TEM images of the myelinated tracts in the dorsal (top rows) and ventral SC (bottom rows). Yellow arrows indicate myelinated axon. Red arrows indicate unmyelinated axon. (**H**) Quantification of dorsal and ventral myelinated axons per area (n = 5 fish).

We further examined the therapeutic effect of citicoline on myelin deficits in *Slc44a1* cKO mice. Citicoline supplementation started from P5 to P21 (**Figure 8**A). Black gold staining revealed a marked reduction in cortical and striatal myelin density in *Slc44a1* cKO mice relative to controls at P21. Interestingly, treatment with citicoline increased the myelin density in *Slc44a1* cKO mice, but not in the controls which was consistent with the results from zebrafish (Figure 8B). Moreover, TEM analysis also revealed that *Slc44a1* cKO mice exhibited a notable increased percentage of myelinated axons in the ON and CC following citicoline treatment (Figure 8C). Taken together, citicoline can effectively ameliorate the myelin deficits caused by *Slc44a1* deficiency.

**Figure 8.**
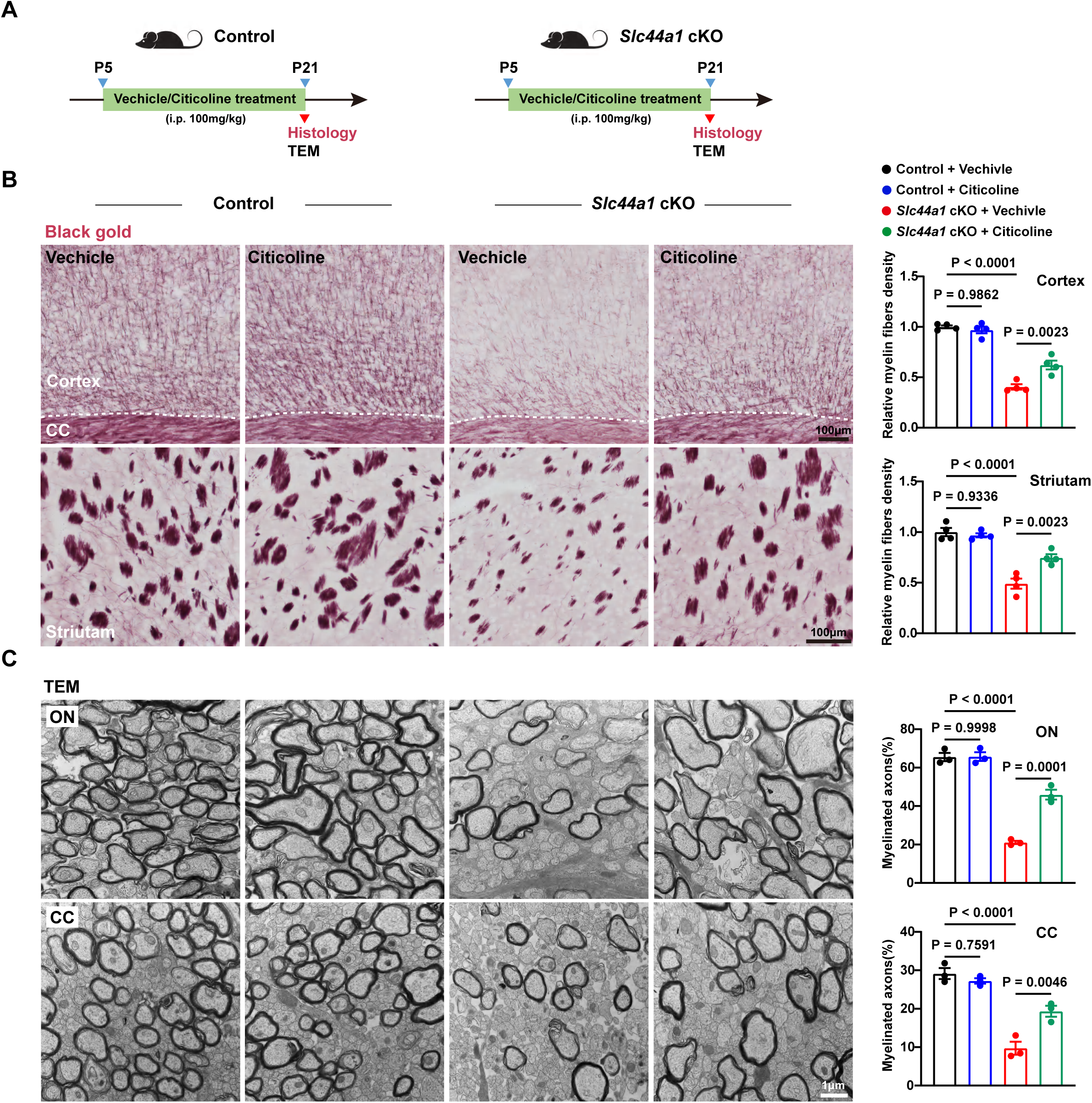
Citicoline supplementation rescues the developmental myelination deficits in *Slc44a1* cKO mice. (**A**) Schematic diagram displaying the time course for citicoline treatment. TEM, transmission electron microscope. (**B**) Black gold myelin staining and quantification of relative myelin fibers density. (**C**) TEM images and quantification of percentage of myelinated axons in the ON and CC.

## 3. Discussion

Choline transporters are categorized into three families according to their affinity for choline: the high affinity choline transporter, SLC5A7, which is unique expressed in cholinergic neurons for the biosynthesis of neurotransmitter acetylcholine;^37^ the low affinity organic cation transporters SLC22A1/2;^38^ and the intermediate affinity choline transporter SLC44A1-5.^39^ Each transporter possesses distinct properties, including choline affinity, tissue distribution, and transport substrate.^40^ SLC44A1 is predominantly expressed in the nervous system.^23,41^ Here, we identified that SLC44A1 was abundantly expressed in mature OLs, but barely detectable in OPCs, neurons, astrocytes and microglia. Its temporal expression was highly correlated with the timeline of brain myelination. Moreover, we found for the first time that conditional ablation of SLC44A1 in OL linage cells leads to significant developmental myelin defects in the CNS of both mice and zebrafish. Consistently, RNA-seq analysis revealed that SLC44A1 knockdown significantly down-regulated gene transcription related to OL maturation and myelination. Importantly, the hypomyelination was identified particularly in the CC, cerebellum, and ON, which is consistent with leukoencephalopathy, cerebellar atrophy, and optic neuritis observed in human patients with SLC44A1 loss by frameshift mutations.^28^ SLC44A1 cKO mice also exhibited a decline in motor coordination and balance, as well as a reduction in cognitive abilities, which aligns with the clinical features of patients.^28^ Myelin formation during development is pivotal for neural integrity and functioning.^42^ Therefore, SLC44A1 deficiency disrupts myelin development in the CNS, which may account for the pathological alterations in cognition and motor skills. It is worth noting that the OL lineage–specific SLC44A1-deficient animals do not recapitulate all the features in the childhood-onset neurodegeneration caused by the systemic loss of SLC44A1. For instance, symptoms such as dysphagia and dysarthria of the patients may be due to the absence of SLC44A1 in muscle cells.^43,44^

In the CNS, myelin formation by OLs along axons represents one of the most remarkable and complex processes in development.^1,2,45,46^ This process entails OPC proliferation, migration, differentiation, and myelinogenesis. Any disruption to this process may result in myelination deficits, potentially leading to neurodevelopmental disorders, such as leukodystrophies.^6,47^ By immunohistochemical staining of CC1 and PDGFRα in CC, which label mature OLs and OPCs respectively, we found that SLC44A1 deficiency decreases the number of mature OLs without affecting OPC populations, indicating a crucial role for SLC44A1 in oligodendroglial maturation. The transparent larval development of zebrafish enables real-time visualization of cellular and molecular processes, providing an excellent model for studying myelination.^48,49^ Long-term time lapse imaging in the SC of zebrafish larvae revealed a significant decrease in both the number and length of myelin segments formed by individual OL in SLC44A1-deficient fish, indicating a reduced myelinating capacity of OLs in SLC44A1 deficiency animals.

The substantial membrane biosynthesis and deposition is required for the metabolically intensive myelinating process of OLs during development.^50^ Myelin is a multilayered stack of uniformly thick membranes containing 70%–80% lipids by dry weight.^50^ The most abundant lipids in myelin are cholesterol, galactocerebroside, sphingomyelin, and PC.^51–56^ Some of these lipid species, such as PC, are direct derivatives of intracellular choline,^14^ whereas the synthesis pathways of others, like galactocerebroside, intersect with choline derivatives.^57^ Here, our RNA-seq data showed that the downregulated genes by SLC44A1 knockdown were significantly enriched in choline metabolic pathway. Among them, PCYT1A, which converts phosphocholine to citicoline and is the rate-limiting enzyme in the biosynthetic pathway of PC, was the most prominently decreased choline metabolic gene. These results indicate that SLC44A1 deficiency leads to defects in the choline metabolic pathway. Our lipidomic data further revealed significant alterations in lipid metabolism (such as cholesterol, fatty acid, glycerolipid) and in lipid composition of myelin sheaths (such as PC and PE) in SLC44A1 knockout animals. Choline-derived lipids are essential structural components and reservoirs of signaling molecules, playing crucial roles to all stages of myelination: from initiation and sheath compaction to long-term maintenance.^57^ Dysregulation of PC and phosphatidylinositols, as well as fatty acid chains length and unsaturation, results in disruptions to myelin structure, membrane stability and curvature, and eventually leads to severe myelination defects.^57^ Our study indicates that SLC44A1 deficiency decreases PC and PE ratio in myelin membrane, in line with the impairment of choline metabolism. One oligodendrocyte, generating 20 to 60 myelin segments and up to 100 turns, has the largest membrane area comparing to any other cells in the body.^58^ Myelin wrapping of single axonal segment takes only a few hours, indicating a high demand for membrane biosynthesis.^59,60^ Therefore, SLC44A1 deficiency limits choline intake and disrupts lipid metabolism and composition of myelin sheaths, ultimately resulting in the deficit of myelin formation during development. Interestingly, fast-growing cells, such as cancer cells, rely on rapid phospholipid synthesis to provide building blocks for their plasma membrane biogenesis and consequently their proliferation. The overexpression of SLC44A1 in various cancer cells may also be linked to the high demand for phospholipids in these cells,^44,61^ implying a possibly shared mechanism by which SLC44A1 supports membrane expansion.

Citicoline serves as a crucial intermediate and rate-limiting step in the biosynthetic pathway of PC.^62^ In this study, we found that citicoline level had the most significant change among the downregulated metabolites in the choline metabolic pathway in OLs following SLC44A1 knockdown. The exogenous delivery of citicoline has shown many beneficial effects, including promoting cell regeneration and survival, restoring lipid balance, and reducing inflammation.^63^ For example, citicoline ameliorated the disease course of experimental autoimmune encephalomyelitis, exerting positive effects on myelin, OLs and axons.^64^ Moreover, citicoline effectively promoted myelin regeneration and reversed deficits in motor coordination following cuprizone-induced demyelination.^64^ In our study, we discovered that citicoline rescues the developmental myelinating defects in SLC44A1 deficiency zebrafish and mice. As an FDA approved drug undergoing clinical trials for the treatment of many neurological disorders, citicoline has shown rare side effects.^65^ Given its strong safety profile and wide availability, citicoline holds promise as a potential therapeutic agent particularly for patients with SLC44A1 deficiency or variation.^26,28^

## 4. Conclusions

In summary, our studies have identified that SLC44A1 is essential for developmental myelination in the CNS of zebrafish and rodents. SLC44A1 deficiency impairs oligodendroglial choline metabolic pathway, downregulates citicoline level, disrupts PC biogenesis and composition of myelin sheath, and eventually leads to deficit of OL maturation and myelinogenesis during development. Citicoline administration ameliorates hypomyelination in SLC44A1 KO animals, presenting a potential strategy for interventions against childhood-onset neurodegeneration of patients with SLC44A1 deficiency or variation (**Figure 9**).

**Figure 9.**
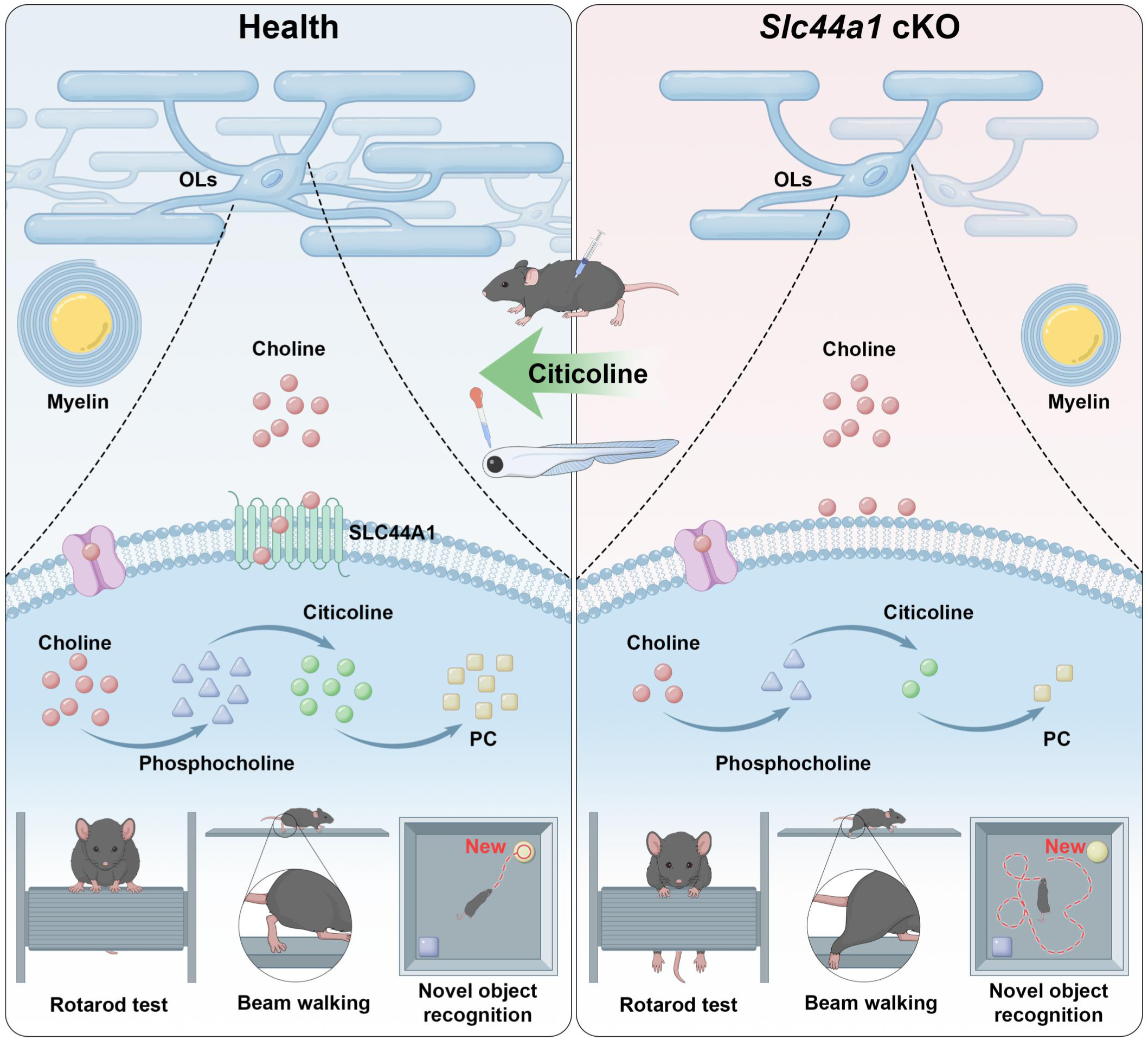
Schematic illustration of effects of SLC44A1 on developmental myelination.

## 5. Experimental Section

### Mice

All animal experiments were performed in adherence with the National Institutes of Health Guidelines on the Use of Laboratory Animals and were approved by the NMU Committee on Animal Care. All mice used in this study were maintained on a C57BL/6 background. *Slc44a1^fl/fl^* mice (ENSMUSG00000028412) were generated by the Shanghai Model Organisms Center, Inc. (Shanghai, China) using CRISPR-Cas9 technology. To conditionally delete *Slc44a1* in OLs, *Slc44a1^fl/fl^* mice were crossed with *Olig1*-Cre mice.^32^ Animals were housed under normal 12-hour light/dark cycle conditions with food and water *ad libitum*.

### Zebrafish Husbandry, Transgenic and Mutant lines

Adult zebrafish (Danio rerio) were husbandry in a recirculating system with a 14 h light/10 h dark cycle at 28±1°C. Freshly fertilized eggs were produced by pairwise mating and raised in dish containing system water.

Tg(Olig2:eGFP) transgenic line was used for labeling OL linage cells.^66^ The transgenic lines Tg(Sox10:mRFP) and Tg(MBPa:eGFP-CAAX) were created by injecting plasmid Sox10:mRFP^67^ and MBPa:eGFP-CAAX,^67^ together with I-SceI enzyme, into one-cell embryos, respectively.

*Slc44a1b*^-/-^ zebrafish were constructed through CRISPR-Cas9 technology,^68,69^ and the following sgRNA sequence were used: CGGTATCACCTGGGCACTGTGG. Cas9 protein (NEB, M0646M) and the sgRNA were microinjected into the animal pole of one-cell stage fish embryos. Injected F0 animals were raised to adulthood and outcrossed to wild-type animals to create F1 offspring. Clutches of F1 offspring were raised to adulthood and genotyped to identify heterozygous carriers of function disrupting mutant alleles.

### Single Oligodendrocyte Labelling

Mosaically labelling OLs was performed as previously reported.^60^ Fertilized eggs were microinjected at the one-cell stage with 1 nL of solution containing 30 ng/μL Sox10:mRFP plasmid DNA, 25 ng/μL Sox10:eGFP plasmid DNA and I-SceI. Animals were screened at 3 dpf.

### Immunofluorescence Staining

The brain tissue or cell slides were fixed with 4% paraformaldehyde (PFA). Subsequently, the samples were blocked with 10% normal donkey serum and permeabilized with 0.2% Triton X-100 in phosphate-buffered saline (PBS) for 1 h at room temperature. Primary antibodies, including anti-Sox10 (1:200, R&D Systems, Catalog #AF2864), anti-Olig2 (1:500; Oasis Biofarm, Catalog #OB-PGP040), anti-PDGFRα (1:150, R&D Systems, Catalog #AF1062), anti-CC1 (1:200, Millipore, Catalog #OP80), anti-SLC44A1 (1:200, Abcam, Catalog #ab110767), anti-MBP (1: 200, Abcam, Catalog #ab7349), anti-PLP (1:200, Oasis Biofarm, Catalog #OB-PRT040), anti-NeuN (1:1,000, MilliporeSigma, Catalog #ABN90), anti-IBA1 (1:500, Synaptic Systems, Catalog #234308), anti-GFAP (1:200, MilliporeSigma, Catalog #G3893), anti-Ki67 (1: 200, Cell Signaling Technology, Catalog #9129), and anti-Cleaved Caspase-3 (1:200, Cell Signaling Technology, Catalog #9661), were diluted in phosphate-buffered saline with 0.2% Triton X-100 and 5% donkey serum. These antibodies were incubated overnight at 4°C, followed by incubation with TRITC-conjugated, FITC-conjugated, or Alexa647-conjugated secondary antibody (1:200, Jackson ImmunoResearch). The samples were then counterstained with Hoechst 33342 (1:1,000, Sigma-Aldrich) for 2 h at room temperature. Fluorescence images were captured using a fluorescence microscope (Dragonfly 200, ANDOR, England) and quantified using ImageJ software (Media Cybernetics).

### In Situ Hybridization

*In situ* hybridization was performed as described previously.^70^ Mice were anesthetized, and brain tissues were isolated and fixed in 4% PFA at 4°C overnight. The tissues were then sequentially transferred into 20% and 30% sucrose solutions for equilibration. Tissues were embedded in optimal cutting temperature compound (OCT) and cryosectioned into 20 µm-thick sections. For *in situ* hybridization (ISH), a *Slc44a1* riboprobe corresponding to nucleotides 2270–3061 of mouse *Slc44a1* mRNA (accession #NM_133891.2, NCBI) was generated in vitro from a cDNA template that had been subcloned into the pEASY®-T&B Zero Cloning vector. Frozen sections were pre-treated and incubated with digoxigenin-labeled nucleic acid probes. The *Mbp*, *Plp1*, and *Pdgfr*α probes were constructed as previous report.^70,71^ Imaging was captured using the Nikon E660FN microscope, and ImageJ software was used for subsequent image analysis.

### Black Gold Myelin Staining

20 µm-thick brain sections were rehydrated in double-distilled water (ddH_2_O) for a duration of 2 minutes (min). Subsequently, the samples were transferred to a preheated gold myelin staining solution (Oasis Biofarm, Catalog #BK-AC001) and incubated at 45°C until the finest myelinated fibers became visible. Following this, the sections were washed twice with ddH_2_O for 2 min each. A sodium thiosulfate solution was then applied, and the sections were incubated at 45°C for 3 min. Afterward, the sections were washed three times with ddH_2_O for 2 min each, and then mounted with Gallic acid mounting medium for examination under a bright-field microscope.

### Western Blot Analysis

Primary cell cultures or tissues were homogenized in RIPA lysis buffer containing a phosphatase and protease inhibitor cocktail on ice (Beyotime, Shanghai, China). The lysate was then centrifuged at 12,000[rpm for 10 min at 4°C. The supernatant was diluted with 5 X loading buffer and boiled at 95°C for 5 min. Proteins were separated using 10-15% polyacrylamide-SDS gels and subsequently transferred to nitrocellulose membranes. After blocking with 10% skim milk for 1 h, the membranes were incubated with the following primary antibodies: anti-SLC44A1 (1:1000, Abcam, Catalog #ab110767), anti-MBP (1:1000, Oasis Biofarm, Catalog #OB-PRB130), anti-PCYT1A (1:500, Santa Cruz Biotechnology, Catalog #sc-376107), anti-β-actin (1:2000, proteintech, Catalog #66009), and anti-GAPDH (1:2000, Proteintech, Catalog #10494-1-AP). The protein bands were analyzed and quantified using Image Lab (ODYSSEY CLX, LI-COR, USA), normalizing the target proteins to the β-actin or GAPDH bands.

### Transmission Electron Microscopy

The optic nerves (ON) and corpus callosum (CC) of mice, as well as the spinal cord (SC) of zebrafish, were isolated and perfused with a phosphate buffer solution containing 2.5% glutaraldehyde and 4% PFA (pH 7.3). The samples were fixed in 1% osmium tetroxide for 45 min, dehydrated, embedded in Araldite resin. Then, the samples were sectioned to a thickness of 70-90 nm using a diamond knife (Leica EM UC7) to 70-90nm, followed by stained with uranyl acetate and lead citrate for contrast enhancement. All the samples were examined using a transmission electron microscope (HITACHI H-7650 or FEI Talos 120) at an accelerating voltage of 100 kV.

### Primary OPCs Culture and Lentiviral Infection

This method has been described previously.^72^ Primary OPCs were isolated from newborn Sprague-Dawley (SD) rats. Mixed cortical glial cell cultures were generated from P0 rats and maintained in Dulbecco’s Modified Eagle’s Medium (DMEM, Invitrogen) supplemented with 10% fetal bovine serum (FBS, Gibco) for 10 days at 37°C under 5% CO_2_. The culture flasks were shaken at 180 rpm for 1 h followed by an additional 18 h at 200 rpm with fresh medium at 37°C. The OPCs were collected from the mixed glia cultures, and the cell suspension was allowed to adhere in uncoated Petri dishes. Isolated OPCs were then plated into culture dishes pre-coated with poly-L-lysine (PLL) and cultured in a proliferation medium supplemented with 0.1% platelet-derived growth factor-AA (PDGF-AA), 2% B27, 1% N2, and 0.1% basic fibroblast growth factor (bFGF). To examine differentiation, OPCs were cultured in Neurobasal medium supplemented with 2% B27 (NB/B27).

For lentiviral construction, the shRNA of *Slc44a1* (Gene ID: NM_001033852.1; TargetSeq: GCTTTCCTAGTCGCTCATT) was constructed by the vector pSLenti-U6 (OBl0, Shanghai, China) and the shRNA with a nontargeting sequence was used as a negative control. The *Slc44a1*-shRNA virus was added to the proliferation medium at the appropriate multiplicity of infection. After an infection period of 8-10 h, the medium was changed to a differentiation medium, and the OPCs were allowed to differentiate for 72 h.

### RNA Isolation and Quantitative PCR (qPCR)

Total RNA was extracted using Trizol (Invitrogen, USA). Each RNA sample was reverse-transcribed into complementary DNA (cDNA) using the Evo M-MLV RT Master Mix (ACCURATE BIOTECHNOLOGY (HUNAN) CO., LTD, ChangSha, China). qPCR was performed on a LightCycler 96 apparatus (Roche) using the SYBR Green Pro Taq HS Premix (ACCURATE BIOTECHNOLOGY (HUNAN) CO., LTD, ChangSha, China). Gene expression was expressed as quantified the mRNA level normalized to that of a standard housekeeping gene (*Gapdh*) using the [[CT method and each reaction was set up with three replicates. Primer pairs were as follows: for *Gapdh*, F: CCATCAACGACCCCTTCATT; R: ATTCTCAGCCTTGACTGTGC; for *Mbp*, F: CATTGTGACACCTCGTACACC; R: GAGCTGGTGTGCTAACTCGT; for *Pcyt1a*, F: CACCGGATTGATTTCGTCGC; R: CTGTGTGGGAGCAAACATGC; *Plp1*, F: TTGGCGACTACAAGACCACC; R: AGCCATACAACAGTCAGGGC; *Cnp*, F: ATGCTGAGCTTGGCGAAGAA; R: GTACAGGTTTGCCCTTCCCA; *Mag*, F: CCTGGATCTGGAGGAGGTGA; R: TCCCATTCACTGTGGGCTTC; *Mog*, F: GTCTATCGGCGAGGGAAAGG; R: AGCATAGGCACAAGGGCAAT.

### BrdU Incorporation Assay

The bromodeoxyuridine (BrdU) incorporation assay was performed according to the previous study.^73^ Briefly, OPCs were plated on PLL-coated glass slides and cultured with a proliferation medium consisting of DMEM/F12 supplemented with 0.1% PDGF-AA, 2% B27, 1% N2 and 0.1% bFGF. BrdU was added to the well and incubated for 6 h. The cell slides were then washed with PBS and fixed in 4% PFA for 10-20 min for subsequent staining. The cell slides were immersed in 2N HCl at 37°C for 30 min, and then washed in 1M sodium borate buffer (pH 8.5) for 10 min. The primary antibody BrdU (1:100, Sigma-Aldrich) was incubated overnight at 4°C.

### RNA-seq Library Construction and Sequencing Analysis

Primary rat OPCs were infected with *Slc44a1*-shRNA lentivirus and corresponding controls for 12 h and then transferred to a differentiation medium. Total RNA was extracted after 3 days of differentiation, with 4 replicate samples in each group, each sample containing 1*10^6^ OLs cells. Total RNA was isolated using Trizol (Invitrogen, USA). Strand-specific libraries were prepared using the TruSeq Stranded Total RNA Sample Preparation Kit (Illumina, USA) according to the manufacturer’s instructions. Briefly, mRNA was enriched using oligo(dT) beads. After purification, the mRNA was fragmented into small pieces using divalent cations at 86°C for 6 min. The cleaved RNA fragments are copied into first strand cDNA using reverse transcriptase and random primers. This is followed by second-strand cDNA synthesis using DNA Polymerase I and RNase H. These cDNA fragments then undergo end repair process, the addition of a single ‘A’ base, and then ligation of the adapters. The products are then purified and enriched by PCR to produce the final cDNA library. Purified libraries were quantified by Qubit 2.0 Fluorometer (Life Technologies, USA) and validated by Agilent 4200 Bioanalyzer (Agilent Technologies, USA) to confirm the insert size and calculate molar concentration. Clusters were generated by cBot with the library diluted to 10 pM and then were sequenced on the Illumina NovaSeq 6000 (Illumina, USA). The library construction and sequencing were performed at Shanghai Biotechnology Corporation.

### Myelin Isolation and Purification

Myelin was isolated by sequential centrifugation on a discontinuous sucrose gradient as described previously.^74,75^ Ultracentrifugation process was performed on a SW41 Ti rotor. Brain tissues were first homogenized in 0.3 M sucrose solution containing 20 mM Tris-Cl buffer (pH 7.45), 2 mM EDTA, 1 mM dithiothreitol (DTT), 100 μM phenylmethylsulfonyl fluoride (PMSF), 10 μg/ml leupeptin, and 10 μg/ml antipain. This homogenate was then layered onto a sucrose density gradient consisting of 0.3 M and 0.83 M sucrose in 20 mM Tris-Cl buffer and 5 mM EDTA at a pH of 7.4, and centrifuged at 75,000g for 35 min. The crude myelin membrane band accumulated at the interface between 0.3 M and 0.83 M sucrose was carefully collected. The sucrose was then removed by washing with Tris-Cl buffer (pH 7.45). The myelin fraction was then subjected to two rounds of hypoosmotic shock and low speed centrifugation at 12,000 g for 15 min to remove cytoplasmic and microsomal impurities. For further purification, the crude myelin was resubmitted to the initial density gradient centrifugation process, along with another cycle of hypoosmotic shock and low-speed centrifugation. To obtain a highly purified myelin fraction, a third density gradient centrifugation was performed. In this step, the myelin was resuspended in 0.83 M sucrose, which was then covered with a layer of 0.30 M sucrose. During centrifugation, the myelin migrated upward and formed a band at the interface between the 0.30 M and 0.83 M sucrose layers. Finally, the purified myelin pellet was resuspended in 1 ml of Tris-Cl buffer and stored at -80 °C.

### Lipidomic Analysis

Lipids were extracted from myelin sheath of mice brain tissue using a modified version of the Bligh and Dyer’s method.^76^ Briefly, 4 mL of chloroform: methanol (2:1) (v/v) was added to samples and incubated at 1,500 rpm for 1 h at 4°C. The samples were then centrifuged and the lower organic phase containing lipids was extracted into a clean tube. Lipid extraction was repeated once by adding 2 mL of chloroform to the remaining aqueous phase, and the lipid extracts were pooled into a single tube and dried in the SpeedVac under OH mode. Samples were stored at -80°C until further analysis. Total protein content was determined from the dried pellet using the Pierce® BCA Protein Assay Kit according to the manufacturer’s protocol.

Lipidomic analyses were conducted at LipidALL Technologies using a Shimadzu Nexera 20-AD HPLC coupled with Sciex QTRAP 6500 PLUS as reported previously.^77^ Separation of individual lipid classes of polar lipids by normal phase (NP)-HPLC was carried out using a TUP-HB silica column (i.d. 150x2.1 mm, 3 µm) with the following conditions: mobile phase A (chloroform: methanol: ammonium hydroxide, 89.5:10:0.5) and mobile phase B (chloroform: methanol: ammonium hydroxide: water, 55:39:0.5:5.5). MRM transitions were set up for comparative analysis of various polar lipids. Individual lipid species were quantified by referencing to spiked internal standards. d9-PC32:0(16:0/16:0), d9-PC36:1p(18:0p/18:1), d7-PE33:1(15:0/18:1), d9-PE36:1p(18:0p/18:1), d31-PS(d31-16:0/18:1), d7-PA33:1(15:0/18:1), d7-PG33:1(15:0/18:1), d7-PI33:1(15:0/18:1), C17-SL, d5-CL72:8(18:2)4, Cer d18:1/15:0-d7, d9-SM d18:1/18:1, C8-GluCer, C8-GalCer, d3-LacCer d18:1/16:0, Gb3 d18:1/17:0, d7-LPC18:1, d7-LPE18:1, C17-LPI, C17-LPA, C17-LPS, C17-LPG, d17:1 Sph, d17:1 S1P, C14-BMP, d3-16:0-carnitine were obtained from Avanti Polar Lipids. GM3-d18:1/18:0-d3 was purchased from Matreya LLC. Free fatty acids were quantitated using d31-16:0 (Sigma-Aldrich) and d8-20:4 (Cayman Chemicals).

### Metabolomic Analysis

This method has been described previously.^78^ Briefly, primary rat OPCs were infected with *Slc44a1*-shRNA lentivirus and corresponding controls for 10 h and then transferred to differentiation medium. Metabolites were extracted after 3 days of differentiation, with 3 replicate samples in each group, each sample containing 5*10^6^ OLs cells. We performed the 600MRM analysis (Biotree, Shanghai, China) with LC-MS/MS. 1500 μL acetonitrile-methanol-H_2_O (2:2:1, containing isotopic internal standards) was added to the Eppendorf tube containing harvested cells. All the Eppendorf tubes were placed in dry ice to pre-cool. The samples were then frozen in liquid nitrogen and were thawed in 37°C water bath. The freeze-thaw cycle was repeated 3 times and the samples were vortexed for 30 s. After sonication in an ice-water bath for 15 min, the samples were incubated at -40°C for 2 h. Then, the samples were centrifuged at 12,000 rpm and 4°C for 15 min. 1,200 μL of supernatant from each sample was transferred to a new Eppendorf tube and was dried with a centrifugal concentrator. 120 μL of 60% acetonitrile was added to the Eppendorf tube to reconstitute the dried samples. The Eppendorf tube was vortexed until the powder was dissolved, followed by centrifugation at 12,000 rpm and 4°C for 15 min. Finally, 60-70 μL of supernatant from each sample was transferred to a glass vial for LC-MS/MS analysis.

The LC separation was carried out using an UPLC system (H-class, Waters) equipped with a Waters Atlantis Premier BEH Z-HILIC column (1.7 µm, 2.1 mm *150 mm). The mobile phase A was a mixture of H_2_O and acetonitrile (8:2) containing 10 mmol/L ammonium acetate, and the mobile phase B was a mixture of H_2_O and acetonitrile (1:9) containing 10 mmol/L ammonium acetate. The mobile phases A and B were adjusted to pH 9 with aqueous ammonia. The autosampler temperature was set at 8°C and the injection volume was 1 μL. An AB Sciex QTrap 6500 plus mass spectrometer was applied for assay development. Typical ion source parameters were as follows: Ion spray voltage: +5000V/-4500V, curtain gas: 35 psi, temperature: 500°C, Ion source gas 1: 50 psi, Ion source gas 2: 50 psi. Raw data files generated by LC-MS/MS were processed using SCIEX Analyst Work Station software (1.7.2) and metabolites quantification was analyzed with BIOTREE Bio Bud (2.1.4).

### Administration of Citicoline

Mice were randomly divided into two groups. The experimental and control groups were administered 100 mg/kg citicoline (MedChemExpress, Catalog #33818-15-4) as previously described^64^ or an equal volume of physiological saline via intraperitoneal injection, respectively. For the zebrafish administration study, embryos were mechanically dechorionated at 1 dpf and the larvae were randomly divided into control and the citicoline-treated groups. They were then transferred into small petri dishes containing with 1 mM citicoline or vehicle. The solutions were renewed every day during the imaging period.

### Open Field Test

The open field test was performed on mice as previously described^79^ with minor modifications. Mice were placed in the corner of an open-field apparatus (40 m × 40 cm) for 10 min of spontaneous exploration with an overhead video-tracking system. The total distance mice traveled, the distance they traveled in the center area (20 cm × 20 cm center square), and the distance they spent and entries in the center area were monitored throughout the experiment. The apparatus was thoroughly cleaned with 75% alcohol between two experiments. The ratio of the distance traveled in the central area to the total distance was used as an indicator of anxiety-related responses.

### Rotarod Test

The rotarod test assesses mouse motor function, balance, and coordination.^80^ Two days before the experiment, the experimental mice were trained 4 times at a speed of 4 rpm for 5 min each, with a 30-min interval between each training session. During the experiment, the rotarod (diameter 3 cm) accelerated from 4 rpm to 40 rpm within 5 min. Mice were placed individually on the rotarod, and a high-definition camera recorded the time the animals spent on the rotarod. The experiment was conducted 4 times, with a 30-min interval between each experiment.

### Beam Walking Test

This test assesses fine motor ability in mice as previously described.^81^ In the modified balance beam walking test, the balance beam (0.5 cm wide) was placed 50 cm above the floor in a dark room, with one end illuminated by a lamp and the other end placed in a box (opaque, 20×10×20 cubic centimeters, containing mouse cage bedding). In the acquisition phase, mice were positioned on the beam at distances of 30, 50, and 70 cm from the illuminated end, and they underwent training for a period of three days. On the test day, high-definition cameras recorded from both sides of the balance beam, recording the frequency of hind limb slips as the mice walked a distance of 80 cm.

### Novel Object Recognition Test

The novel object recognition test was performed as previously described.^82^ The test is divided into a training phase and a testing phase, with a high-definition camera recording the mouse’s activity. During the training phase, mice were placed in the experimental apparatus without any objects to adapt for 5 min. Then, each mouse was individually placed into the arena containing two identical objects, equidistant from each other, and allowed to explore the objects for 10 min. In the testing phase, which began 5 min after training, one object from the training phase and one novel object was placed either side of the walls of the apparatus at a distance of 5 cm (in the same positions as in the testing phase, with the left and right positions randomized). Mice were placed in the experimental apparatus with a familiar object and a novel object and explored for 10 min. The recognition index (time spent exploring the new object/total time spent exploring both objects) was used as an indicator of recognition memory.

### Statistical Analysis

Data analysis in this study was performed using GraphPad Prism software. Data were analyzed using one-way ANOVA with Tukey post hoc test or Dunnett’s multiple comparisons for multiple groups, and unpaired Student’s t-test was used for two groups. The data are presented as mean ± SEM. The value of P < 0.05 was considered statistically significant.

### Ethics Statement

The animal study was reviewed and approved by the Institutional Animal Care and Use Committees of NMU and to relevant guidelines and regulations.

## Supporting Information

Supporting Information is available from the Wiley Online Library or from the author.

## Acknowledgments

We thank Dr B Appel at University of Colorado School of Medicine for providing the Tg(Olig2:eGFP) transgenic line and the Sox10:mRFP plasmid; Dr Jiulin Du at Center for Excellence in Brain Science and Intelligence Technology, Chinese Academy of Sciences for WT/AB line; Dr Q. Richard Lu at Cincinnati Children’s Hospital Medical Center of Cincinnati for providing the *Olig1*-Cre mice; Dr Yun Li at Tongji University School of Medicine for providing IBA1 antibody; and Dr Masato Inazu at Institute of Medical Science, Tokyo Medical University for providing the information of SLC44A1 antibody. This work was supported by grants to CH from Ministry of Science and Technology of the People’s Republic of China STI2030-Major Projects 2022ZD0204700, to PL from National Natural Science Foundation (31400920), and to ZL from National Natural Science Foundation (82401564).

## Author contributions

Q.C., X.C., and Z.L. contribute equally to this work. Q.C., P.L., and C.H. designed and performed research. X.C. performed bioinformatic analysis. Q.C., Z.L. and Q.S. carried out most of the experiments and data processing. W.Z. provided clinical information support. H.H. and M.Q. provided *in situ* hybridization support. Q.C., Z.L. and P.L. wrote the manuscript. Z.S., M.Q., and C.H. revised the manuscript.

## Data availability

The data sets used and/or analysed during the current study are available from the corresponding author on reasonable request.

---

**Figure S1.**
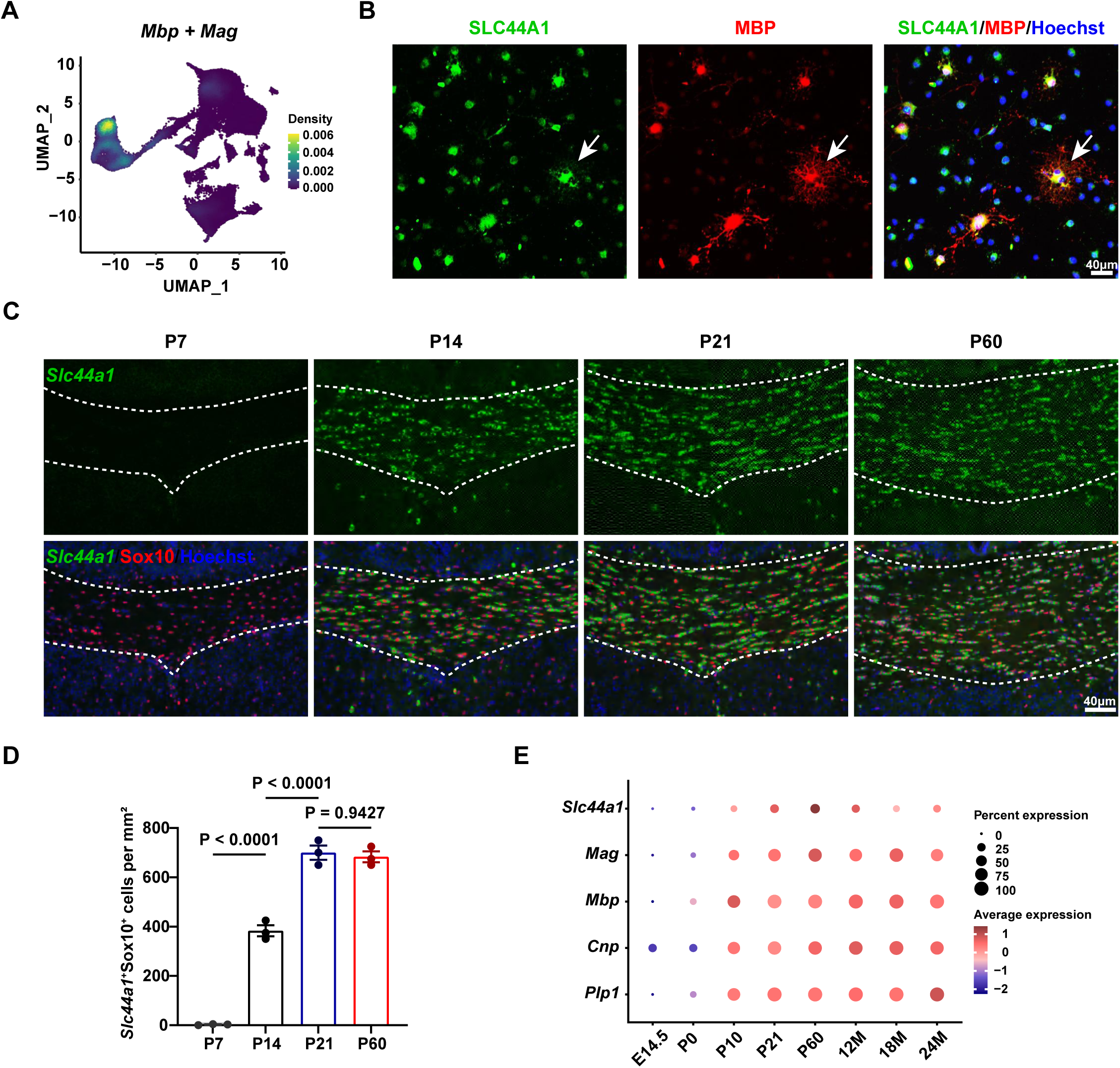
Dynamic expression of SLC44A1 in OLs during development. (**A**) UMAP plots showing the expression of *Mbp* and *Mag* mRNA. (**B**) Immunostaining of SLC44A1 and MBP in primary rat OLs (n = 3 independent experiments). (**C**) *In situ* hybridization for *Slc44a1* (green) and Immunostaining for Sox10 (red) in the CC of wildtype mice at the indicated time points (n = 3 mice). (**D**) Quantification of *Slc44a1*^+^Sox10^+^ cells in the CC of wildtype mice at the indicated time points (n = 3 mice). (**E**) Dot plots showing the expression of *Slc44a1*, *Mag*, *Mbp*, *Cnp*, and *Plp1* mRNA levels at the indicated time points.

**Figure S2.**
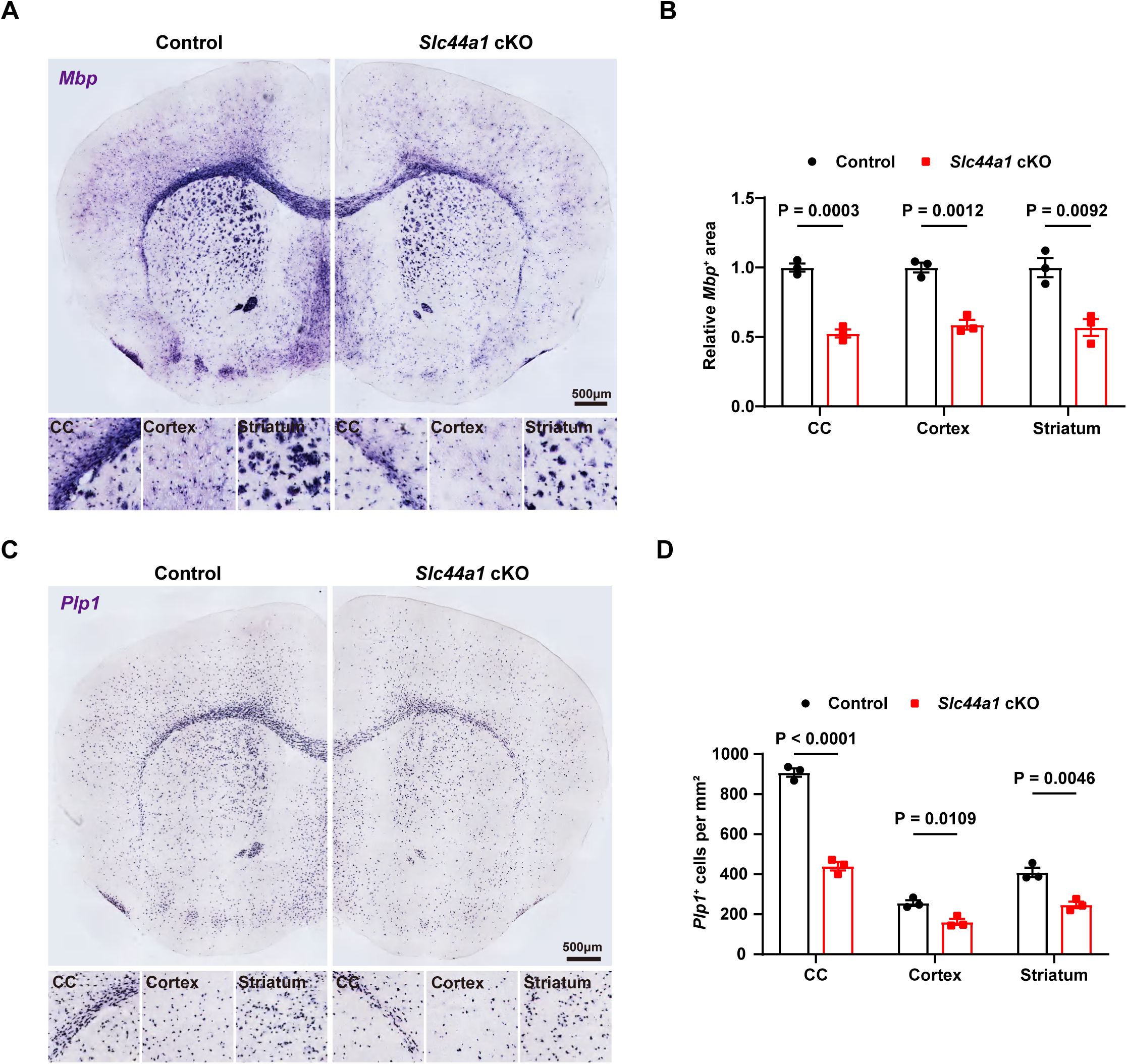
SLC44A1 is required for proper CNS myelination. (**A**) *In situ* hybridization for *Mbp* in brains of control and *Slc44a1* cKO mice at P21. (**B**) Quantification of relative *Mbp^+^* area in the CC, cortex, and striatum of control and *Slc44a1* cKO at P21 (n = 3 mice). (**C**) *In situ* hybridization for *Plp1* in brains of control and *Slc44a1* cKO mice at P21. (**D**) Quantification of *Plp1*^+^ cell per mm^2^ in the CC, cortex, and striatum of control and *Slc44a1* cKO at P21 (n = 3 mice).

**Figure S3.**
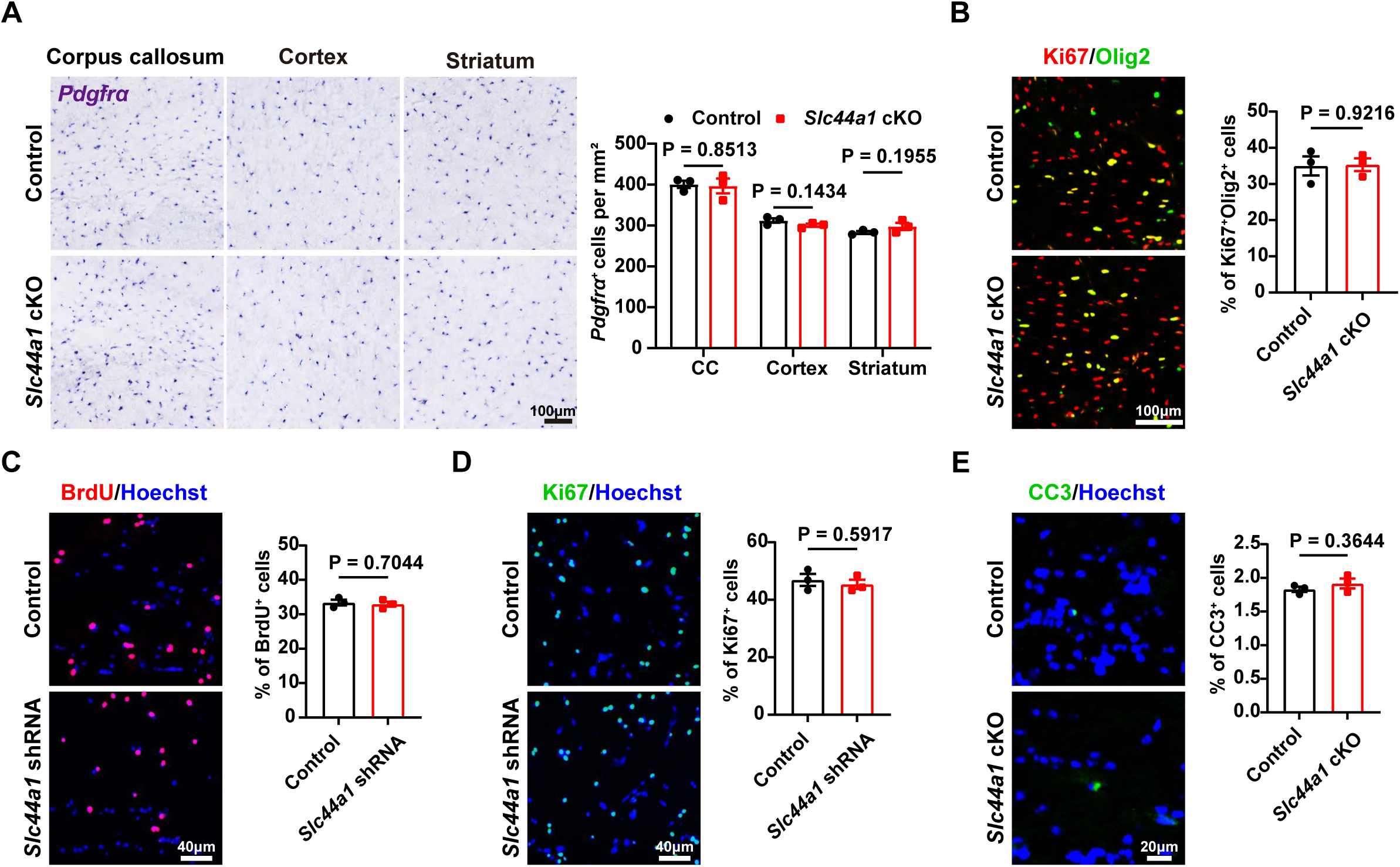
SLC44A1 deletion in the OLs does not affect OPC proliferation. (**A**) Left, *In situ* hybridization for *Pdgfr*α in the CC, cortex, and striatum of control and *Slc44a1* cKO mice at P21. Right, quantification of *Pdgfr*α^+^ cells (n = 3 mice). (**B**) Left, immunostaining for Ki67 and Olig2 in the CC of control and *Slc44a1* cKO mice at P21. Right, quantification of percentage of Ki67^+^Olig2^+^ cells among total Olig2^+^ cells (n = 3 mice). (**C**) Left, immunostaining for BrdU in control and *Slc44a1* shRNA-treated OLs. Right, quantification of percentage of BrdU^+^ cells (n = 3 independent experiments). (**D**) Left, immunostaining for Ki67 in control and *Slc44a1* shRNA-treated OLs. Right, quantification of percentage of Ki67^+^ cells (n = 3 independent experiments). (**E**) Left, immunostaining for cleaved caspase-3 (CC3) in the CC of control and *Slc44a1* cKO mice at P21. Right, quantification of percentage of CC3^+^cells (n = 3 mice).

**Figure S4.**
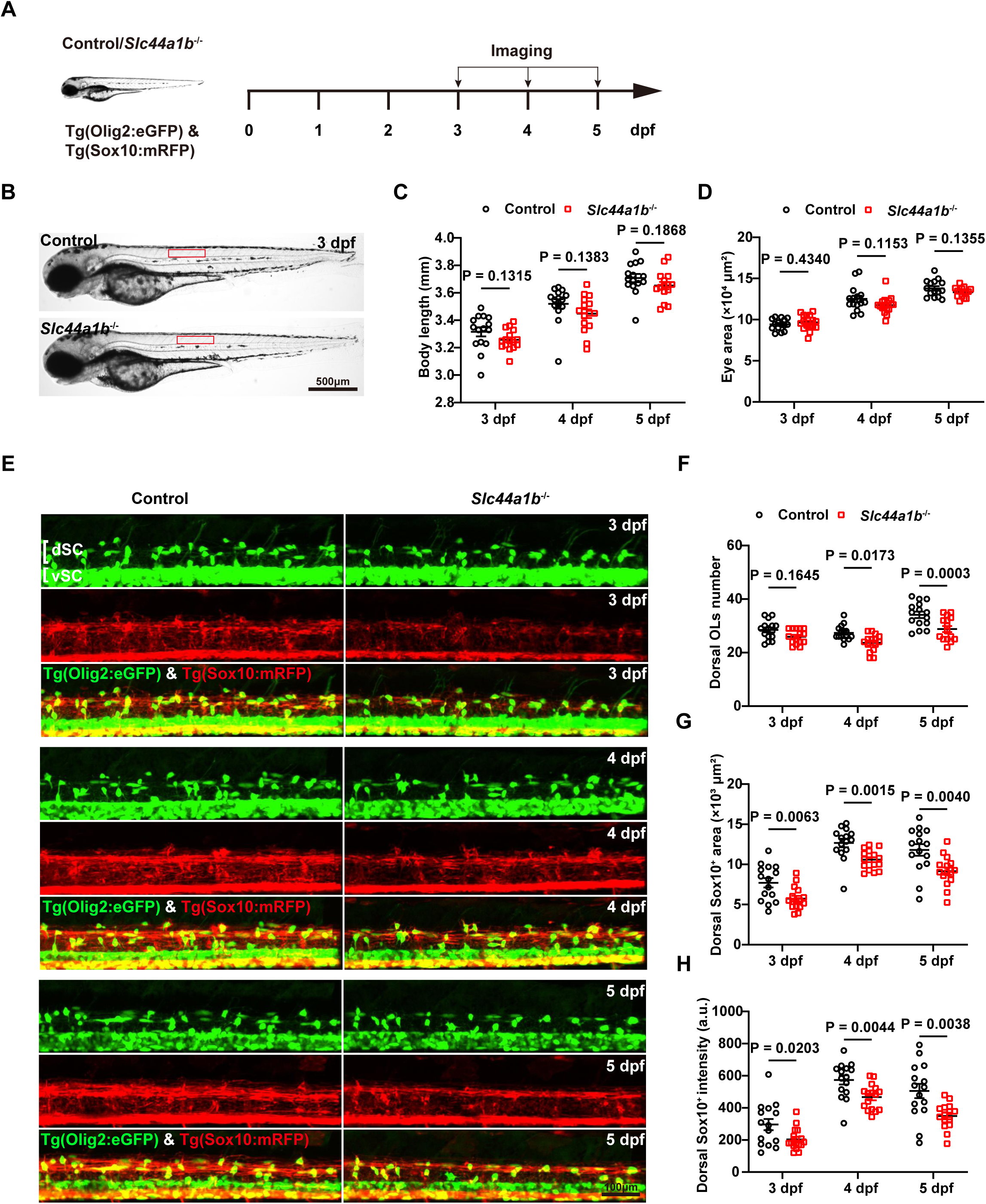
*Slc44a1b* deletion leads to impairments in OL maturation and myelination. (**A**) Diagram of the timeline of the experiment. (**B**) Bright field imaging shows the normal morphology and the area (9 - 12 somites) being analyzed in (**E**). (**C**, **D**) Measurements of larval length and eye area (n ≥ 14 fish). (**E**)The fluorescent images of Tg(Olig2:eGFP) and Tg(Sox10:mRFP) transgenic lines in control and *Slc44a1b*^-/-^ larvae. (**F**) Quantification of dorsal OLs number (n ≥ 14 fish). (**G**, **H**) Quantification of dorsal RFP area and intensity.

**Figure S5.**
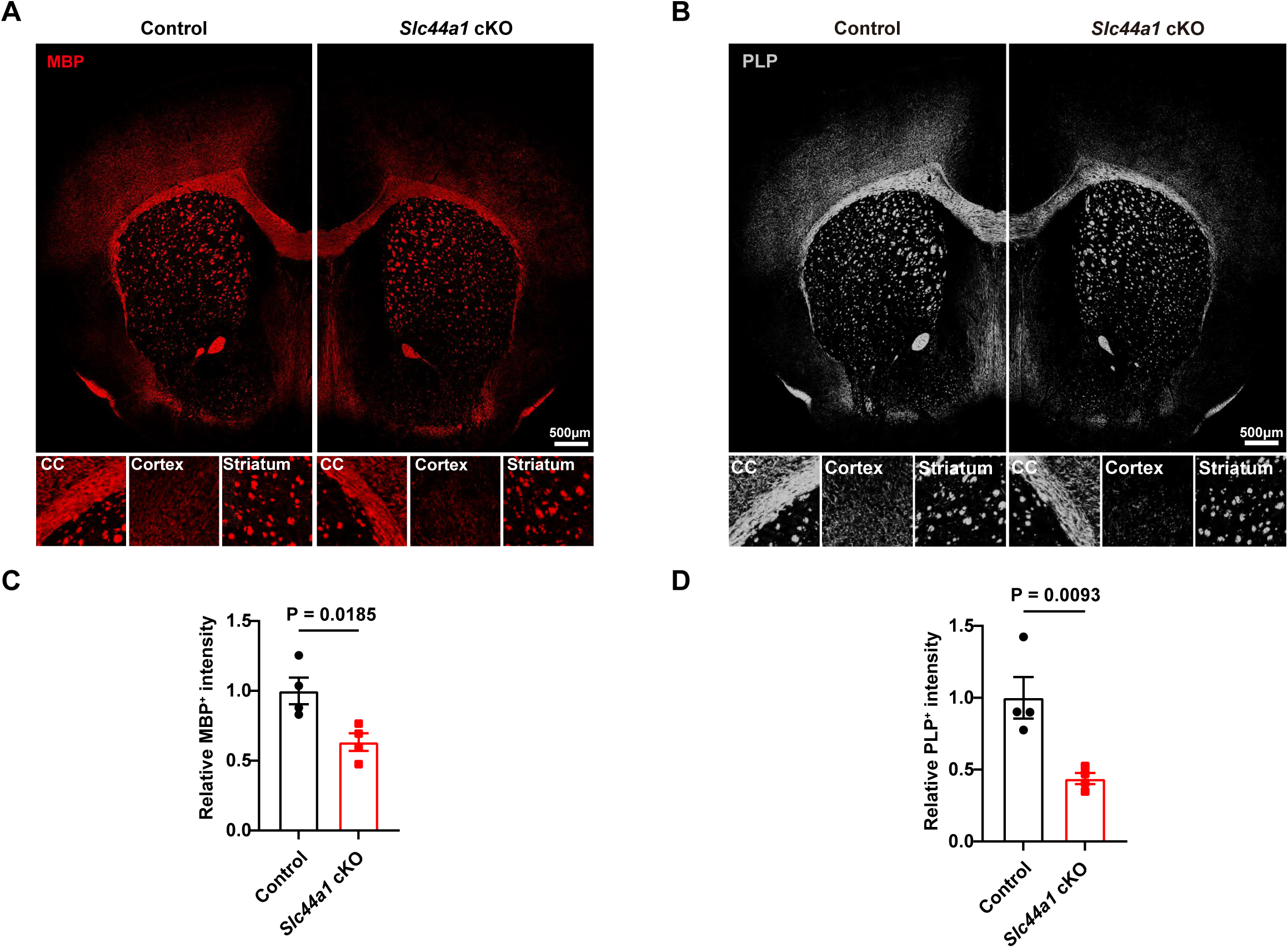
Adult *Slc44a1* cKO mice exhibits myelination deficits. (**A**) Immunostaining for MBP in multiple brain regions of control and *Slc44a1* cKO mice at P60. (**B**) Immunostaining for PLP in multiple brain regions of control and *Slc44a1* cKO mice at P60. (**C**) Quantification of relative MBP^+^ intensity in the whole brain of control and *Slc44a1* cKO mice at P60 (n = 4 mice). (**D**) Quantification of relative PLP^+^ intensity in the whole brain of control and *Slc44a1* cKO mice at P60 (n = 4 mice).

